# Mbd4 and MutSα protect cells from spontaneous deamination of 5-methylcytosine

**DOI:** 10.1101/2024.12.17.628571

**Authors:** Rebecca A. Bilardi, Christoffer Flensburg, Zhen Xu, Emily B. Derrick, Andrew Kueh, Ian J Majewski

## Abstract

5-Methylcytosine (5mC) is a common source of somatic mutations. Deamination of 5mC to thymine generates a G/T mismatch, which occurs spontaneously and must be repaired prior to DNA replication to avoid mutation. We generated genetically engineered mice and cell lines to define DNA repair pathways that protect against methylation damage. We observed a low background mutation rate in mouse bone marrow or colon, typically 0.2-0.5 CG>TG mutations/genome/day. This increased 3-7 fold in cells lacking the glycosylase Methyl-binding domain 4 (Mbd4), one of the few glycosylases capable of excising thymine from G/T mismatches. We found no role for Thymine DNA glycosylase (Tdg) in methylation damage repair. Instead, our results support cooperation between Mbd4 and the mismatch repair (MMR) complex MutSα (Msh6:Msh2), evident through elevated rates of methylation damage in Msh6-deficient cells; increasing to 2.6-4.8 CG>TG mutations/genome/day in primary cells and up to 13.9 CG>TG mutations/genome/day in cell lines. Our findings support the view that MutSα has DNA repair activity outside of replication. While loss of Mbd4 elevates methylation damage selectively, the broader functionality of MutSα explains why mutational signatures linked to Msh6-deficiency are variable and reflect the replicative history of the cell.

## Introduction

The genome is under constant attack from endogenous and exogenous sources of damage. A complex network of DNA repair pathways exists within cells to detect and correct DNA damage (1). DNA repair pathways operate with astounding efficiency, but they are not infallible. This means somatic mutations accumulate as we age and contribute to the genesis of cancers and other diseases. One of the major genomic signatures of ageing comes from CG>TG transitions that occur at sites of methylated cytosine due to spontaneous deamination; these mutations are evident in normal cells (2), in cancers (3) and in the germline (4). Deamination of 5-methylcytosine (5mC) results in a thymine that is mismatched with guanine, this will be referred to as methylation damage. Methylation damage is responsible for the gradual depletion of CG sites and this has had a major influence on the organisation of the genome. Although specific repair mechanisms have been described to counter methylation damage, they are insufficiently understood.

The Catalogue Of Somatic Mutations In Cancer (COSMIC) has been used to define common mutational signatures (3,5). COSMIC Single Base Substitution signature 1 (SBS1) is characterised by CG>TG transitions and is regularly used to quantify methylation damage. Genomic studies have revealed that SBS1 accumulates with age, in both normal tissues and in cancers, and it is detected across a wide variety of species (6). We previously detailed a highly specific accumulation of methylation damage in individuals with biallelic loss of the DNA glycosylase Methyl-binding domain 4 (*MBD4)*. As a result, these individuals are predisposed to colorectal polyposis, acute myeloid leukaemia, uveal melanoma and other cancers (7–10). MBD4 is one of many substrate specific glycosylases that catalyse the first step in base excision repair (BER). Following methylation damage, MBD4 excises the mispaired thymine and allows replacement of the cytosine (11–13). Looking across a range of cancers and experimental models, we found that loss of MBD4 resulted in the accumulation of CG>TG mutations at a broadly consistent rate, of between 1-2 mutations per genome per day (7,9). The rate of spontaneous deamination of methycytosine in naked DNA *in vitro* has been determined to be 5.8 × 10^-13^/second (14). If the *in vivo* rate is comparable, it would equate to ∼2-3 events per genome per day, suggesting that other DNA repair mechanisms contribute to the repair of methylation damage.

Thymine DNA glycosylase (TDG) is another glycosylase in the BER pathway and based on its *in vitro* substrate specificity it is thought to target G/T mismatches and products of active cytosine demethylation, including formyl and carboxyl cytosine (15,16). Genetic analysis of Tdg in mice is complicated by early embryonic lethality (17), consistent with an important regulatory role during development. Inactivation of TDG did not increase the mutation burden or reveal an obvious DNA repair defect in a human induced pluripotent stem cell model, albeit in an *in vitro* system and over a relatively short culture period (18). The precise contribution of TDG to BER, and to the repair of methylation damage remains unclear.

Mismatch repair (MMR) is a replication linked pathway that detects and repairs mismatches that occur during DNA replication (19). MMR has been hypothesized to have replication independent functions, and recent studies suggest this includes elimination of methylation damage. Fang et al. found a marked accumulation of CG>TG mutations in some human cancers that carry inactivating mutations in the MMR components MSH2 or MSH6 (20). MSH2 and MSH6 form a heterodimer, called MutSα, which can bind mismatches or small (1-2 nucleotide) insertions or deletions. Human MSH6 was originally isolated as a G/T mismatch binding protein (21). The link between MutSα and methylation damage is further supported by genomic profiling of tissue biopsies from individuals with constitutional mismatch repair deficiency (CMMRD). Sanders et al. showed that biallelic loss of MSH2 or MSH6 was associated with elevated levels of methylation damage in primary tissues and cancers (22). This increase seemed to be specific to MutSα, because it was not evident in individuals with inherited defects in MLH1 or PMS2, which partner to form MutLα. This is unexpected, as MutSα and MutL cooperate to mediate recruitment of downstream factors that coordinate strand excision and resynthesis. It remains unclear whether MutSα works in partnership with other DNA repair pathways to recognise methylation damage and mediate repair.

The purpose of this study was to investigate the key DNA repair pathways that coordinate the response to methylation damage (Figure 1A) and to reveal any cooperativity between them. To do this we have looked at the rate and spectrum of mutations in primary cells from mice lacking *Mbd4*, *Tdg* or *Msh6*, either alone or in combination. We went on to disrupt these pathways in cell line models and in malignant tissues to assess how somatic mutation profiles change in highly replicative tissues and in response to transformation. Our results confirm that Mbd4 and Msh6 contribute to methylation damage repair *in vivo*, and are consistent with a broader role for MutSα outside of its contribution during DNA replication.

**Figure 1:**
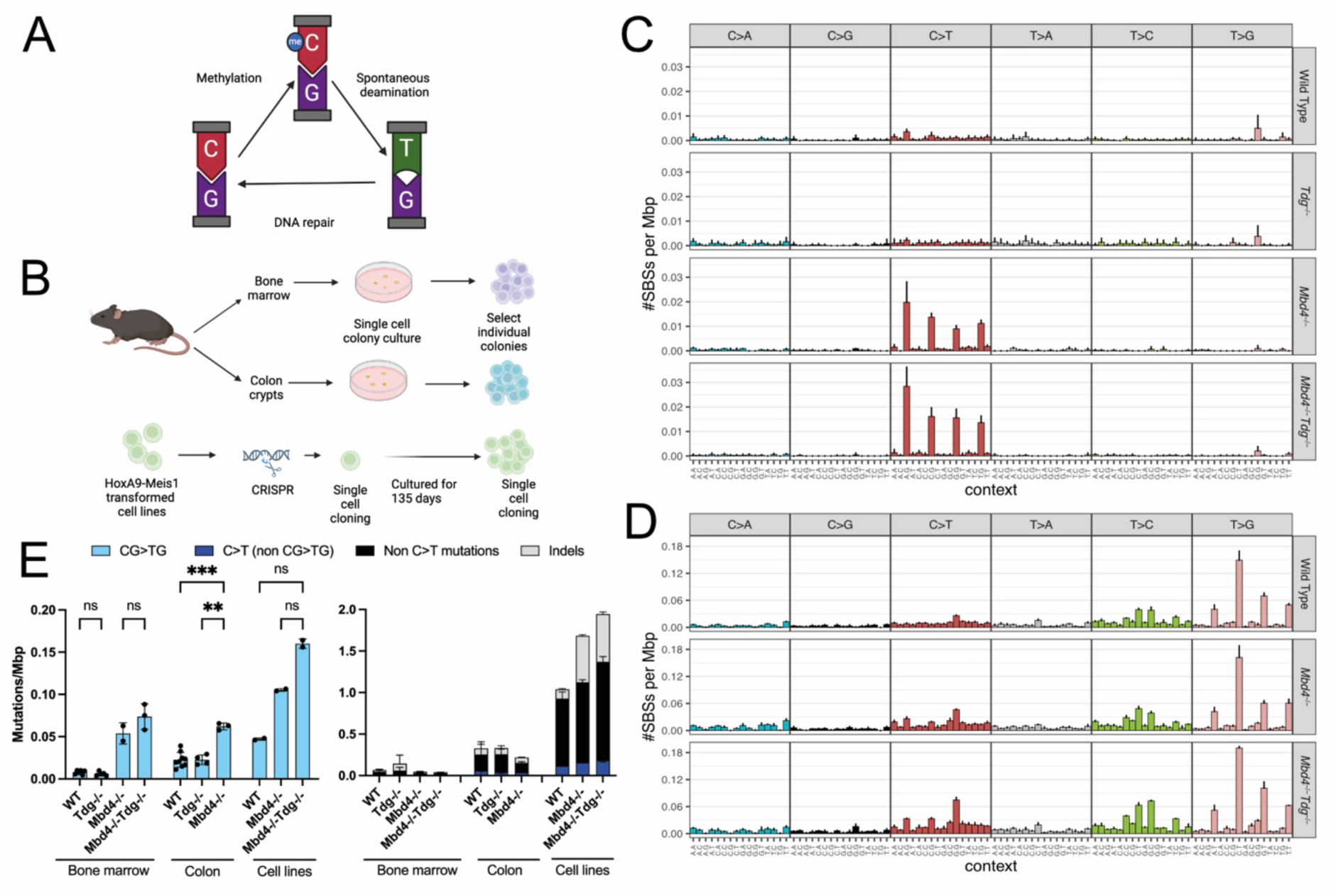
Assessing the role of Mbd4 and Tdg in methylation damage repair. (A) Schematic showing the cycle of cytosine methylation by DNA methytransferase enzymes to 5-methylcytosine (5mC), spontaneous deamination to thymine and its repair. Created in BioRender. Bilardi, R. (2024) https://BioRender.com/f76z261 (B) Diagram outlining the various methods used to study clonal primary cells (bone marrow and colon crypt) and transformed cell lines (HoxA9-Meis1). Created in BioRender. Bilardi, R. (2024) https://BioRender.com/d41y371 (C) Mutational signatures derived from genomes from 3-4 month old WT (N=6), *Tdg^−/−^* (N=5), *Mbd4^−/−^* (N=2) and *Mbd4^−/−^Tdg^−/−^* (N=3) bone marrow colonies. (D) Mutational signatures from WT (N=2), *Mbd4^−/−^* (N=2) and *Mbd4^−/−^Tdg^−/−^*(N=2) HoxA9-Meis1 cell lines, where N = number of clones per genotype. (E) CG>TG mutations per Mbp across the different genotypes and tissues (at left). Individual points represent data from separate colonies. Stacked bar chart of other relevant mutation types in the same samples (at right). All data in this figure represents the mean and SD. Significance tested in Prism with ANOVA with Welch’s correction, select comparisons are shown, asterisks denote statistical significance (*** p<0.001) and non-significant comparisons are marked ns.

## Methods

### Animal models and ethics

All mouse experiments and breeding were approved by the Walter and Eliza Hall Institute Animal Ethics Committee (2017.003, 2018.046, 2020.021 and 2021.048). The following mouse strains were obtained from Jackson Laboratories: B6.SJL-Ptprc^a^ Pepc^b^/BoyJ (JAX stock #002014, referred to as Ly5.1), and B6.Cg-*Mbd4^tm1Wed^*/J (13) (JAX stock #004989, referred to as Mbd4) was maintained by regular backcrossing to C57BL/6 mice. Conditional knockout alleles were generated using CRISPR-mediated editing to introduce loxP sites to flank exon 3 of *Tdg* or to flank exon 2 of *Msh6*. The alleles were designed so that Cre-mediated recombination disrupts the reading frame. Heterozygous floxed mice were crossed with constitutively active CMV-Cre mice to generate knockout alleles (referred to as *Tdg* and *Msh6*). The Tdg flox mice were crossed to Vav-iCre (23) (JAX stock # 008610) to generate haemopoietic specific deletion (referred to as Vav Tdg) or to Rosa26 CreERT2 (24) (JAX stock # 008463) to generate mice with tamoxifen inducible, whole body deletion (referred to as CT2 Tdg). Mbd4 and Msh6 lines were intercrossed to generate double mutants (referred to as Mbd4 Msh6). Mbd4 and Msh6 colonies were maintained by heterozygous intercrosses to avoid accumulation of mutations in the germline. Knockout animals were not used for breeding. Genotyping primers and PCR conditions are outlined in Table S1.

### Tamoxifen treatment

Mice were treated with 5mg tamoxifen dissolved in corn oil via oral gavage for two consecutive days to induce gene deletion in Tdg CT2 mice. For these experiments, mice in the control arm also received tamoxifen treatment.

### 5-fluorouracil administration

5-fluorouracil (5FU) (Sandoz) was diluted in PBS and mice were injected intravenously once with 150mg/kg. Mice were bled twice weekly (∼30μl total volume per bleed) via the submandibular vein and blood counts monitored by diluting 1/10 prior to automated blood analysis.

### Automated blood analysis

Either whole or diluted blood was analysed using an ADVIA 2120i Haematology system using CBC/Diff settings for mouse.

### Bone marrow colony culture and sequencing

Bone marrow from 3- to 6-month-old mice was harvested and seeded in 35mm petri dishes at 10,000 cells/plate in Dulbecco’s Modified Eagles’ Medium with 20% foetal bovine serum, 0.3% BactoAgar, 100 ng/ml mSCF (WEHI), 2U/ml hEPO (Eprex, Janssen Australia), and 10 ng/mL mIL-3 (WEHI). After 8-10 days growth at 37 degrees, 10% CO_2_ in a humidified atmosphere, individual colonies were isolated using a fine glass pipette and a dissecting microscope. DNA was isolated using Qiagen QIAmp Micro DNA kit. After quality assessment on an Agilent Tapestation 4200 using genomic DNA reagents, DNA was sheared to 300-400bp for library preparation using a Covaris S220 sonicator. Libraries were prepared with Illumina TruSeq Nano DNA kit and sequenced with 150bp PE reads at ∼300M reads per sample using an Illumina NovaSeq 6000 (AGRF, Parkville, Australia) or BGI DNBSEQ G-400 sequencer (BGI, Hong Kong). Library quantification and pooling was performed by the sequencing service provider.

### Colon colony culture and sequencing

Colon organoid culture was based on a published protocol with minor modifications (25). The distal colon (two thirds total) from 3 month old mice, or mice 3 months post tamoxifen treatment in the case of Tdg CT2, was rinsed in PBS, briefly sterilised in a weak (0.04%) sodium hypochlorite solution and then incubated in 3mM EDTA in order to liberate crypts from the intestinal wall. The crypts were then treated with dispase (Invitrogen) and tryPLE (Gibco) and filtered to obtain a single cell suspension. Single cells were plated in 50% Matrigel (Corning LifeSciences) domes with culture media consisting of 70% mouse IntestiCult (Stem cell tech), 30% YH2CM (26), 10 µM Y27632 (Sigma-Aldrich) and 1x penicillin/streptomycin (Gibco). Incubation was at 37°C in a humidified environment with 10% CO_2_. Individual colonies were picked after 8-9 days and rinsed in ice cold PBS, to remove residual matrigel. Cells were lysed and DNA isolated using Qiagen QIAmp Micro DNA kit. DNA was also extracted from Matrigel for genome sequencing, to enable bioinformatic filtering of variants from any contaminating DNA. Libraries were prepared with Illumina TruSeq Nano DNA kit and sequenced with 150bp PE reads at ∼300M reads per sample using an Illumina NovaSeq sequencer.

### CRISPR cell line knockouts

Transformed myeloid cell lines were generated by infecting foetal liver cells with HoxA9-Meis1, as previously described (27). Routine culture was in DMEM, with addition of 10% foetal bovine serum (FBS) and 10 ng/ml IL-3 (PeproTech) at 37°C in a humidified atmosphere containing 10% CO_2_. Cell lines were generated from wild type, *Mbd4^−/−^* and *Tdg^−/−^* foetal liver cells (from *Vav Tdg* mice). Cell lines were transduced with FUCas9Cherry (Addgene #70182, a gift from Marco Herold) to drive expression of Cas9-mCherry and enable CRISPR/Cas9 editing (28). Single guide RNA (sgRNA) sequences targeting mouse *Msh6* (TTGGCAAAGGCGCTCCAAAG) and *Mbd4* (GATTTTACTGTACTGCCGAA) were cloned into pKLV-U6gRNA (BbsI)-PGKpuro2ABFP (Addgene #50946). Cells were transduced with sgRNAs to inactivate *Msh6* in wild type and *Mbd4^−/−^* cell lines, to generate *Msh6^−/−^*and *Mbd4^−/−^Msh6^−/−^* derivates. *Mbd4^−/−^Tdg^−/−^*cells were generated by transducing *Vav Tdg^fl/−^* cells with sgRNAs targeting *Mbd4.* Single cell cloning was performed and genomic DNA isolated to permit genotyping around the PAM site. PCR primers and genotypes are provided in Table S2.

Amplicons were sequenced on an Illumina MiSeq (San Diego, CA, USA) (28). Cells with two distinct edits were selected to lessen the potential for selection of clones with chromosomal loss. For long-term cultures, cells were grown in regular culture for 135 days, before a second round of single cell cloning. Genomic DNA was extracted from two independent descendent clones for each genotype, together with the ancestral clone (collected at day 0) using the DNeasy Blood and Tissue Kit (Qiagen). Whole genome sequencing was performed as a service by BGI using DNBseq PE150.

### Generating mouse embryonic fibroblast (MEF) cell lines

Timed mouse matings were used to obtain embryos at e13.5-14.5 or e10.5 in the case of *Tdg^−/−^*. Pieces of the embryo were mechanically disrupted to produce a single cell suspension in PBS, spun down and resuspended in DMEM with 10% FBS and allowed to adhere and grow in cell culture for 2 passages. Cells were then transfected with pSG5 SV40 large T antigen plasmid (a gift from David Huang) with FuGeneHD (Promega). Cells were maintained in culture until their growth was stable and consistent; at this point they were considered transformed. All cell lines maintained in culture were tested monthly for mycoplasma.

### Western blotting

Whole cell lysates were typically obtained by suspending cell pellets or mechanically disrupting mouse tissues in RIPA buffer containing Protease Inhibitor Cocktail (Roche) and 1 mM PMSF (Sigma Aldrich) and incubating on ice for 30 min. The samples were centrifuged at 10,000g for 20 min, the supernatant containing soluble proteins was removed and the protein concentration determined via Bradford assay (Bio-rad). Protein concentration was normalised between samples and run on 4-12% bis-tris SDS-PAGE gels (Invitrogen). In some cases, cell pellets were lysed directly by boiling in sample buffer to avoid proteolytic degradation. Gels were blotted onto nitrocellulose membranes using the iBlot system (Invitrogen), blocked in 5% skim milk in PBS-T (PBS with 0.1% Tween-20) for 30 min at room temperature and then probed with the following primary and secondary antibodies: anti-Mbd4 (Santa Cruz Biotech D-6 or R&D Systems AF5935), anti-Msh6 (AbCam EPR3945), anti-Tdg (clone 9B3, derived at WEHI by immunising a mouse with recombinant human TDG glycosylase domain, aa 111-348), anti-Mlh1 (Novus biologicals SP08-04), anti-Msh2 (Cell Signalling D24B5), anti-Pms2 (Novus biologicals 163C1251), anti-Hsp70 (clone N6), anti-rabbit HRP (Southern Biotech 4030-05), anti-mouse HRP (Southern Biotech 1070-05), and anti-goat HRP (Southern Biotech 4010-05). Immobilon Forte Chemiluminescent reagent (Millipore) was added to membranes prior to imaging with a Biorad ChemiDoc XRS+ system.

### Tumour transplantation

Spontaneously arising thymomas from *Msh6^−/−^* or *Mbd4^−/−^Msh6^−/−^*mice were cryopreserved as single cell suspensions in foetal bovine serum with 10% DMSO and later thawed and transplanted into Ly5.1 mice. For the grafts, recipient mice were injected intravenously with 1 × 10^6^ live thymoma cells. Mice were sacrificed at the ethical endpoint when tumour burden was impacting health. The timeframe for this was 16-26 days post-transplant. At sacrifice, spleens were prepared into a single cell suspension and stained for Ly5.1 (clone A20) and Ly5.2 (clone S450-15-2). Viable Ly5.2 single positive cells were sorted and DNA extracted. Bulk bone marrow from the primary mouse was used as a source of germline control DNA. Whole genome library preparation and 150bp PE sequencing was performed as a service by BGI using DNBSEQ G-400.

### Bioinformatics

#### Sequence alignment and variant calling

Sequencing reads were aligned to mm10 with bwa (v0.7.17-r1188) (29), deduplicated using Picard toolkit (30) and analysed with superFreq (v1.4.3) (31). To improve the accuracy of mutational signatures and mutation rates (mutations/Mbp), we censored repeat regions (Simple_repeat or Low_complexity in repeatMasker (32)). Samples were dynamically assigned a read depth cut, where most samples (63 out of 78) were assigned a minimum read depth of 15, but for some samples the read depth cut was decreased to consider at least 40% of the non-repeat genome, down to a minimum cut of 10 (7 samples) (Table S5). The median callable region, bases outside of repeat regions with read depth at or over the read depth cut, across all samples was 2.19 Gbp (range 0.23 Gbp to 2.25 Gbp). As our samples were predominantly clones of single cells with minimal copy number alterations, we expect variant allele frequencies (VAFs) of somatic variants to be distributed around 0.5. Filters were applied requiring VAF ≥0.3 with a minimum of 4 variant supporting reads. Inside the callable region, we analysed variants with a superFreq somaticP score >0.1, and removed variants detected in any other sample from the same batch. Further, variants were filtered to remove any positions where a Matrigel variant was called.

For driver mutation detection, we used more permissive settings and considered all mutations with at least 2 variant reads and no quality flag from superFreq (which assesses base and mapping quality metrics), but otherwise placed no requirements on somaticP, VAF or read depth. Variants were filtered against the matrigel genome as above. Variant annotation was performed with the R package VariantAnnotation (33) and mutations were reported for mouse homologs of COSMIC census genes (34).

#### Assessment of mutational signatures

Standard mutational signatures – single base substitutions (SBS) in a trimer context – were calculated with MutationalPatterns (35). Insertions and deletions (indels) were included as 8 classes of indels based on their size (1, 2-3, 4-6 and 7 or more bases). This resulted in an SBS/indel signature that considered 104 variant classes. For decomposition into mismatch repair signatures 1-6 (MMR-1 to 6) (22), we considered the 96 SBS trimers, along with the total summed number of insertions and deletions separately (a total of 98 variant classes).

Variants were annotated against replication strand using replication fork direction data (36), where variants were classified as forward or reverse based on the strand of the C in a mutated C:G base pair, and based on the T in a mutated T:A base pair. To provide greater discriminatory power, variants in a region with replication fork direction bias smaller than 0.3 were removed from the analysis.

Variants were annotated for replication timing based on their location in the genome using data from mouse endodermal cells as reference (37). We then assessed the relative abundance of mutations found in regions that are replicated early (first 30%) compared to those replicated late (latest 30%). CG methylation status was determined using whole genome bisulphite sequencing data from bone marrow (38), using only sites with read depth of at least 5. Sites were classified as methylated if ≥50% of reads indicated the presence of 5mC.

#### Variant simulation as controls

We simulated SBSs for each sample to serve as a control for testing bias in replication strand and timing. We excluded the repeatMasker regions above, and randomly generated 10 times the SBSs of the original sample within each trimer context to preserve the mutational signature in the simulated variants.

#### Comparison of mutation rates in NCG trimers

To more accurately compare mutation rates between each NCG trimer, mutation frequencies were adjusted to account for trimer frequency in the non-repeat genome. We considered all bases outside of the filtered repeatMasker region in mm10 and hg38 for mouse and human genomes, while the calculation for callable regions for exomes considered all Ensembl exons, padded by 100 bases on either side. All NCG mutation counts were corrected to mm10 genome trimer frequency, before re-normalising to fractions across the four NCG trimers. We made use of several reference sets of somatic mutations, including those in mouse colonic crypts (6), normal human colon (39), *MBD4*-deficient AML (7), *MSH6*-deficient cell lines (18), and tumours selected from the Cancer Genome Atlas (TCGA) that exhibited biallelic loss of *MSH6* and were enriched for methylation damage (at least 80% CG>TG mutations).

## Results

### Tdg does not contribute significantly to the repair of methylation damage

A range of biochemical studies suggest that TDG has the capacity to excise thymine (40,41), but this has not been verified *in vivo*. To address this, we developed a new targeted allele of *Tdg* to enable conditional deletion of exon 3 (Figure S1A) with Cre recombinase. A B6 deleter strain (CMV-Cre) was used to generate a constitutive knockout allele. Timed matings with *Tdg^+/−^* mice revealed that *Tdg^−/−^* animals died at approximately e11 (Table S3), consistent with earlier studies that identified an important role during embryogenesis (17). *Tdg* deletion by Tamoxifen treatment of CT2 Tdg mice at 7 weeks had no obvious impact on the overall health of the mice, similarly to a previous report (42). Deletion of *Tdg* did not impact blood counts measured at 6 weeks (Figure S1B-D), or recovery from bone marrow stress (Figure S1E-G). Loss of Tdg protein was confirmed in two separate conditional models; using myeloid cell lines and thymocytes derived from Vav Tdg mice, where Cre is expressed in the blood, and colon cells from CT2 Tdg mice, where Cre-ER is expressed broadly and activated by treatment with tamoxifen (Figure S1H-I).

To assess the contribution of Tdg to the repair of methylation damage, we performed mutational profiling using clonally derived cells, either with loss of *Tdg*, *Mbd4* or both genes. This analysis was performed with primary cells from bone marrow and colon from 3-4 month old mice, as well as in HoxA9-Meis1 transformed myeloid cell lines (Figure 1B). Loss of Mbd4 resulted in a specific increase in methylation damage, in both the bone marrow progenitors (0.054±0.13 CG>TG/Mbp, compared to 0.008±0.002/Mbp in the wild type) and colon organoids (0.062±0.004 CG>TG/Mbp, compared to 0.023±0.009/Mbp in the wild type) (Figure 1C and E). This equated to an average of 1.1±0.26 CG>TG mutations/genome/day in *Mbd4^−/−^* bone marrow and 1.46±0.09 mutations/genome/day in colon, which agreed with our earlier observations (7,9). In contrast, loss of Tdg did not raise mutation numbers above wild type in these tissues (Figure 1C, E and Figure S2). In the *Tdg^−/−^* bone marrow progenitors, the methylation damage rate was 0.006±0.002 CG>TG/Mbp, which increased to 0.023±0.06 CG>TG/Mbp in the colon organoids, but these levels were consistent with the wild type (Figure 1C and E and Figure S2). Combined loss of *Mbd4* and *Tdg* resulted in a similar number of mutations to loss of *Mbd4* alone in bone marrow, 0.074±0.015 CG>TG/Mbp (Figure 1C and E), suggesting no major compensatory role for Tdg. We observed similar changes in CG>TG mutation burden in HoxA9-Meis1 transformed myeloid cell lines in which genetic approaches and CRISPR editing were used to inactivate *Tdg* and *Mbd4* (Figure 1D). The HoxA9-Meis1 cell lines were grown for 135 days before single cell cloning to allow for sufficient DNA damage to accumulate. The overall mutation rate was higher in the cell lines, compared to primary tissues, with evidence of an established background signature (27). The cells with combined loss of *Tdg* and *Mbd4* exhibited slightly higher level of CG>TG mutations, but this was not significantly greater than the *Mbd4* single knockout and occurred in the context of a higher overall mutation rate (Figure 1E). Therefore, contrary to *in vitro* studies, our results suggest that Tdg does not contribute to the repair of methylation damage in primary mouse tissues or cell lines.

### Assessing the contribution of Msh6 to the repair of methylation damage

The MutSα complex has been implicated in the repair of spontaneous methylation damage. To assess the relative contribution of MutSα, we generated an *Msh6* knockout mouse model. First we used CRISPR to mediate insertion of loxP sites flanking exon 2 of *Msh6* (Figure S3A). A null allele was generated by crossing to a CMV-Cre strain. *Msh6^+/−^*mice were intercrossed to generate homozygous knockout mice and wild type littermates. *Msh6^−/−^* mice were born healthy and within expected frequencies (360 offspring analysed by Chi-square test, p = 0.0794). Monitoring of peripheral blood revealed no major anomalies in young adult mice (Figure S3C). We examined the stability of Msh6 and other MMR components in the Msh6 knockout model. Western blotting confirmed loss of Msh6 protein in *Msh6^−/−^* cells across various tissues (Figure 2A and Figure S3B). Loss of Msh6 resulted in destabilisation of Msh2 in lysates from primary thymocytes (Figure 2A) and MEFs (Figure S3D). MSH6 and MSH2 levels are also well correlated in quantitative proteomics data from human cancer cell lines (43) (Figure S4). There are prior reports of direct protein-protein interaction between MBD4 and MMR pathway components (44), including evidence that MBD4 may stabilise MLH1 (45). However, we found no evidence that loss of Mbd4 impacts the stability of MMR components in primary thymocytes (Figure 2A) or MEFs (Figure S3D). Nor was there evidence of correlation between the level of MBD4 and MMR components in cancer cell lines (Figure S4).

**Figure 2:**
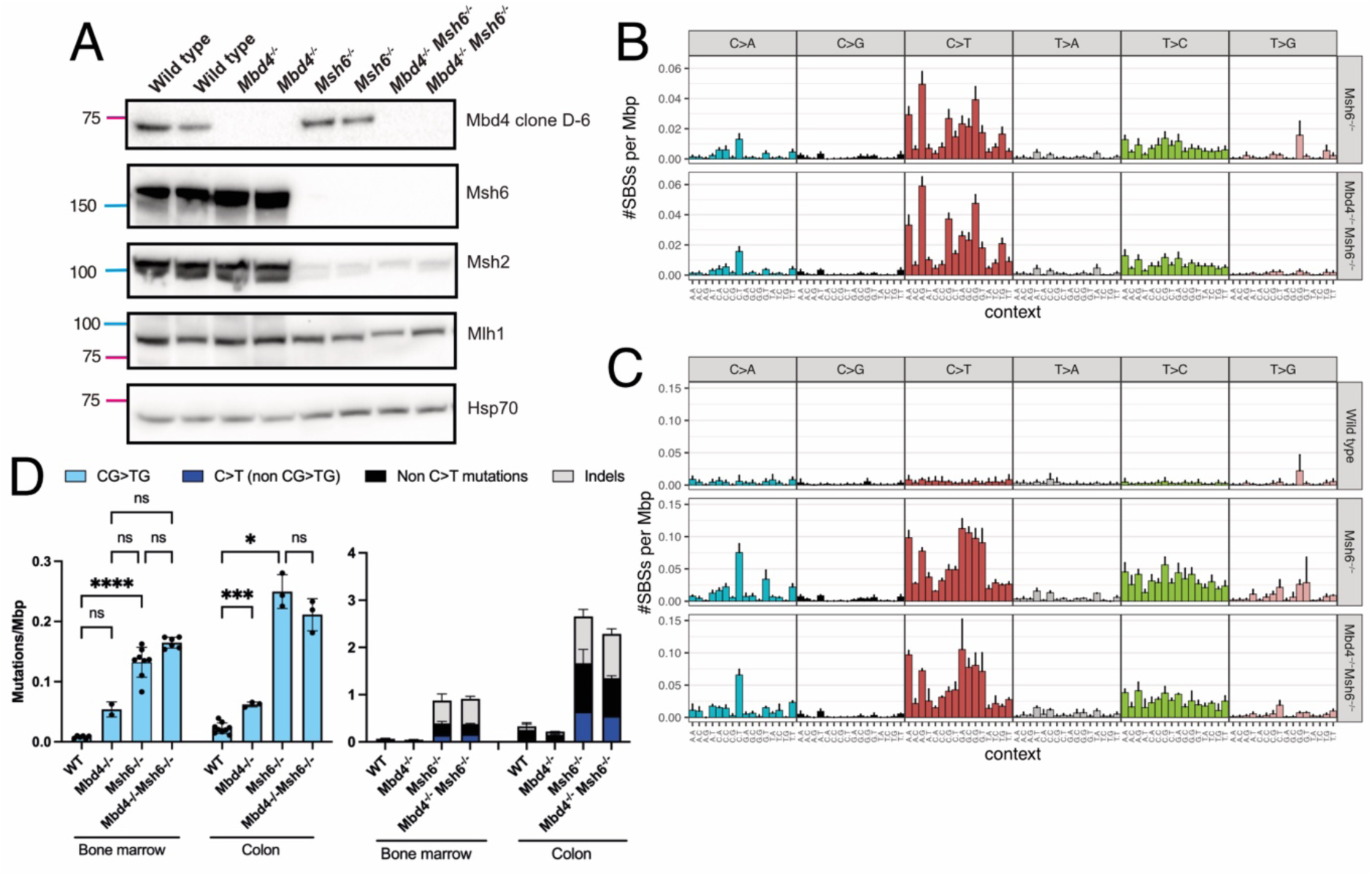
Mutational signature profiling in *Msh6* knockout mice. (A) Western blot of thymocytes from young adult mice (10-18 weeks old). Molecular weights are shown at left (kDA). Two samples were assessed per genotype. Anti-Mbd4, Santa Cruz Biotech clone D-6. (B) Mutational signatures derived from genomes from 3-4 month old *Msh6^−/−^* (N=8) and *Mbd4^−/−^Msh6^−/−^*(N=6) bone marrow colonies and (C) WT (N=9), *Msh6^−/−^* (N=3) and *Mbd4^−/−^ Msh6^−/−^* (N=3) colon colonies. Error bars show SD (D) CG>TG mutations per Mbp across the different genotypes and tissues (at left) and stacked bar chart of other relevant mutation types in the same samples (at right). D includes WT and *Mbd4^−/−^* data presented in Figure 1 for comparison. Bar plots show the mean and SD. Significance tested in Prism with ANOVA with Welch’s correction, selected comparisons are shown, asterisks denote statistical significance (* p<0.05, ****p<0.0001) and non-significant comparisons are marked ns.

We sought to define mutational signatures linked to Msh6 loss in primary mouse tissues, either alone or combined with loss of Mbd4. Genome sequencing was performed for clonal cultures from bone marrow or colon from 3-4 month old *Msh6^−/−^* and compound *Mbd4^−/−^Msh6^−/−^*mice. We found that loss of Msh6 was associated with emergence of a classic mismatch repair deficient (MMRd) signature, including both single base substitutions (SBS) and small insertions and deletions (indels). There was a marked accumulation of SBS in bone marrow progenitors (0.523±0.081 SBS/Mbp), which was predominantly C>T and T>C transitions, and small indels (0.484±0.141 indels/Mbp) (Figure 2B and D). The SBS fraction included a significant amount of methylation damage (0.132±0.025 CG>TG/Mbp, or 2.65±0.5 CG>TG/genome/day), roughly a 2.5-fold increase over the rate seen in *Mbd4^−/−^* bone marrow. Combined loss of Mbd4 and Msh6 resulted in a similar mutation profile as loss of Msh6 alone, and despite a small increase in overall mutation numbers, there was no significant difference in the rate of total SBS (p>0.999) or CG>TG mutations (p=0.129). This pattern was generally consistent using organoids grown from colon crypts from 3-4 month old mice, but with some tissue-specific differences (Figure 2C and D and Table S4). The overall mutation rate was approximately 2-3 fold higher in colon compared to bone marrow (comparing within each genotype), and the Msh6-deficient profiles contained even more C>A and non-CG C>T mutations. This increase in classic MMRd mutations may reflect differences in replicative history between colon and bone marrow. Loss of Mbd4 or Msh6 in the colon was associated with an increased rate of methylation damage, 0.062±0.004 and 0.25±0.028 CG>TG/Mbp, respectively, when compared to the wild type (0.023±0.009 CG>TG/Mbp). Combined loss of Mbd4 and Msh6 produced a similar level of CG>TG mutations (0.211±0.027 CG>TG/Mbp) to loss of Msh6 alone (one way ANOVA with Welch’s correction p=0.823, Figure 2D).

### Assessment of mutational signatures in mouse models of malignancy

Mutational signature analysis in human tumours with loss of MSH6 (20,22) indicates a degree of variability, likely reflecting differences in the timing of the onset of MMRd, the replicative history and potentially cell type specific factors. Mice with loss of Msh6 are also cancer prone (46). In our strain, which was generated on a C57BL/6 background, the *Msh6^−/−^* mice had a median survival of 9 months (Figure 3A). The mice developed a range of malignancies including lymphoma, myelodysplasia, intestinal tumours and other solid cancers. Mice with combined loss of Mbd4 and Msh6 exhibited a similar range of malignancies and had a median survival of 8 months, which was not significantly different to loss of Msh6 alone (Log-rank (Mantel-Cox) p=0.968, Figure 3A). This mirrors earlier work that found loss of Mbd4 did not accelerate tumourigenesis in mice lacking Msh2 or Mlh1 (47). We sought to determine the impact of transformation on mutational signatures in the haemopoetic compartment. First, we performed mutation profiling on thymomas that arose spontaneously in *Msh6^−/−^* (n=3) and *Mbd4^−/−^Msh6^−/−^* animals (n=2). The thymomas were transplanted into Ly5.1 recipient mice to expand the cell population, and flow sorted to obtain a high leukaemic cell fraction. Somatic mutations were detected in expected driver genes including multiple independent hits in *Notch1*, *Ikzf1*, *Trp53,* and *Pten* (Figure S5). The thymomas displayed a characteristic MMRd signatures with a high mutation burden, with both genotypes averaging ∼14 mutations/Mbp (Figure 3B and D). Combined loss of Mbd4 and Msh6 produced a similar pattern of somatic mutations to loss of Msh6 alone, with no significant differences between the groups. In a second approach, we used CRISPR to inactivate *Mbd4* and *Msh6* in Hoxa9-Meis1 transformed myeloid cells and performed long-term cultures to allow DNA damage to accumulate. Genome sequencing of the resulting clones revealed an MMRd signature in both the *Msh6^−/−^* and *Mbd4^−/−^Msh6^−/−^* cell lines (Figure 3C and D and Figure S6). The total mutation burden in the cell lines was much higher than in the primary tissues, with rates increasing to 21.3 mutations/Mbp in the *Msh6^−/−^*cells and 17.1 mutations/Mbp in the double mutant. There was no significant difference in the levels of methylation damage between the *Msh6^−/−^* and *Mbd4^−/−^Msh6^−/−^* cell lines (Figure 3D, p=0.45).

**Figure 3:**
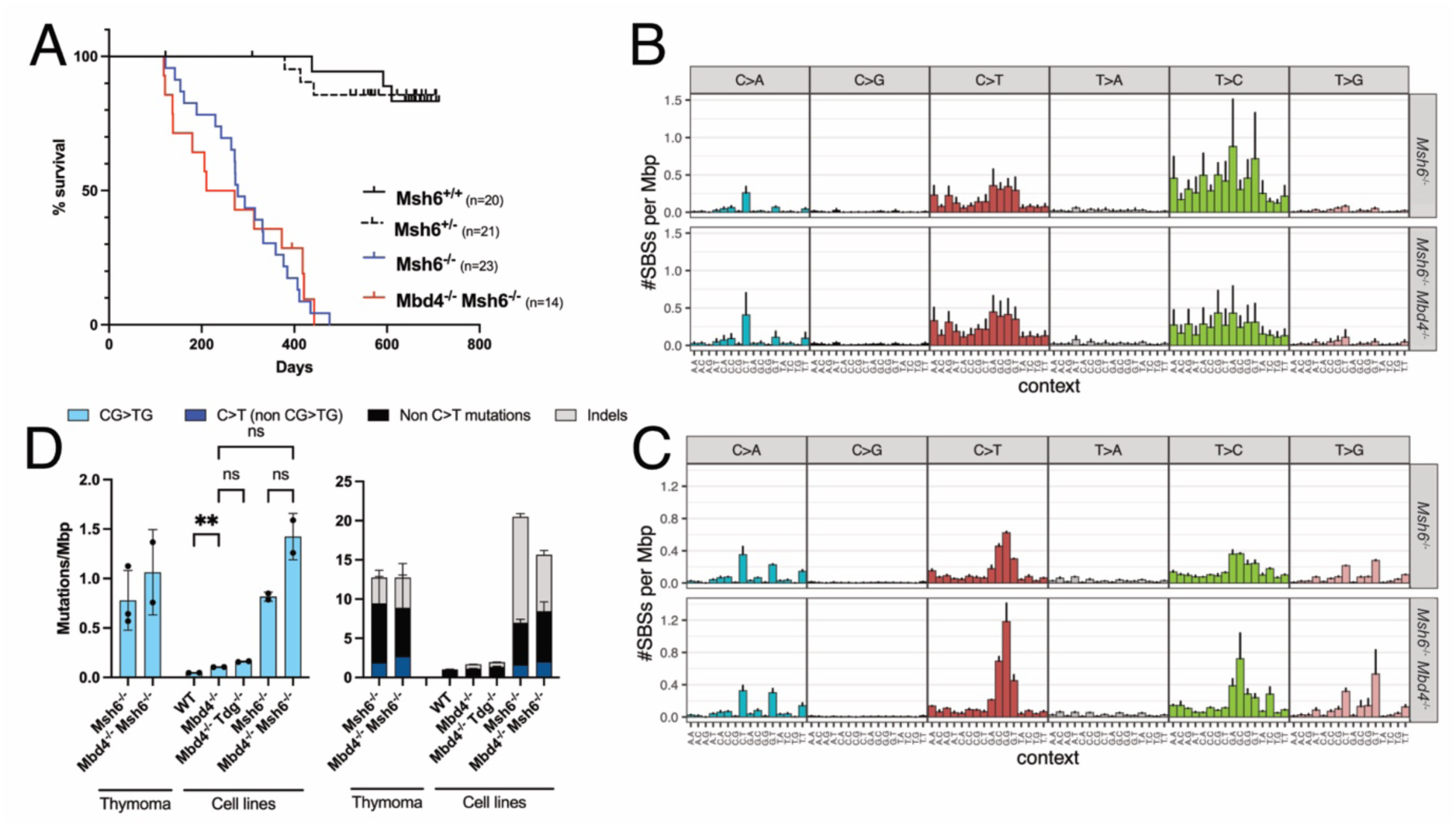
Survival analysis for *Msh6* knockout mice and genomic profiling of transformed cells. (A) Overall survival of *Msh6^−/−^*and *Mbd4^−/−^Msh6^−/−^* mice compared to heterozygotes and wild type controls. Mice were housed in the same facility. Mice were euthanised when they showed signs of ill-health. Mice were censored when euthanised for non-tumour related causes (e.g. fighting wounds) or end of experiment. (B) Mutation signatures derived from genomes from thymomas from *Msh6^−/−^*(N=3) and *Mbd4^−/−^Msh6^−/−^* (N=2) mice, and (C) *Msh6^−/−^*(N=2) and *Mbd4^−/−^Msh6^−/−^* (N=2) myeloid cell lines, where N = independent mice with thymomas or clones per genotype for cell lines. All myeloid cell lines were single cell cloned. (D) CG>TG mutations per Mbp across the different genotypes and tissues (at left) and stacked bar charts of other relevant mutation types in the same samples (at right). Data shown in B-D represent mean and SD. Significance tested in panel D via Prism with ANOVA with Welch’s correction, selected comparisons are shown.

### Comparative analysis of mutational signatures and methylation damage across different experimental systems

We sought to compare the mutation profiles derived from our mouse models to those derived from primary tissues and cancers from people with CMMRD (22). Sanders et al found that methylation damage was enriched in CMMRD cases with loss of MutSα (MSH6 or MSH2), which resulted in a much higher ratio of SBS to indels. While this ratio was roughly 1:1 for loss of most MMR components, it was ∼5:1 in MSH6-deficient individuals. For Msh6-deficient primary samples from mouse models we found roughly equivalent numbers of substitutions and indels, with a ratio close to 1:1 in bone marrow and 2:1 in colon. Indels were rare in wild type control tissues and Mbd4-deficient samples, leading to higher ratios (Figure 4A). Additionally, the representation of 1bp deletions tended to be higher than that of 1bp insertions in CMMRD patients with loss of MutSα (either MSH2 or MSH6), whereas loss of MutL components was associated with equivalent numbers, or favoured insertions. We confirmed a skew towards deletions in both the *Msh6^−/−^* and *Mbd4^−/−^Msh6^−/−^* bone marrow progenitors (Figure 4B).

**Figure 4:**
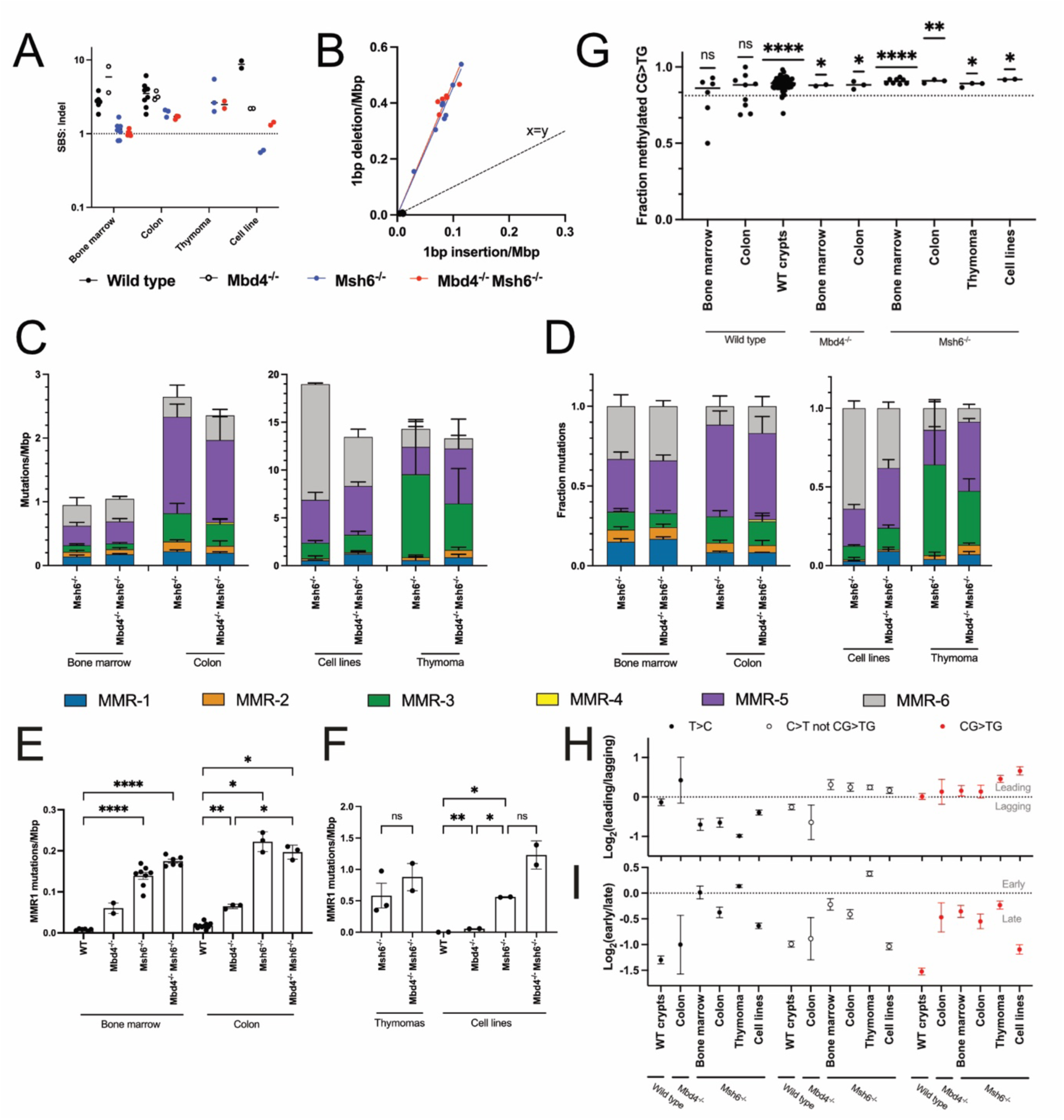
MMRd signature deconvolution and genomic context of mutations. (A) Ratio of SBS to indels in 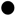 Wild type, 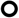 *Mbd4^−/−^,* 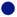 *Msh6^−/−^*or 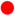 *Mbd4^−/−^Msh6^−/−^* samples separated by tissue. Each point represents an individual sample. (B) The ratio of the number of 1bp insertions to 1bp deletions was calculated for bone marrow colonies for each genotype. Each point represents an individual colony and a regression line is fit with a forced 0 intercept, a control line is set to represent a 1:1 ratio. Deconvolution of mutations into the 6 MMR signatures identified in Sanders *et al*., 2021, shown as a rate (C) or as a proportion (D). MMR1 mutation rates across (E) primary and (F) transformed cells. Significance tested in Prism with ANOVA with Welch’s correction, select comparisons are shown, asterisks denote statistical significance (* p<0.05, ** p<0.01, *** p<0.001, **** p<0.0001) and non-significant comparisons are marked ns. (G) The fraction of CG>TG mutations occurring at methylated sites. The dotted line represents the average methylation rate for CG sites across the dataset. Significance was tested using a one sample t and Wilcoxon test. Logit transformed (H) replication strand or (I) replication timing status of various mutation types compared to random simulations via the following formula Log_2_((observed mutations leading/observed mutations lagging)/(simulated mutations leading/simulated mutations lagging)). Error bars represent standard deviation, using poisson distributions around the number of mutations.

To better separate classical MMRd damage from methylation damage, we used signature deconvolution to quantify each process. We used a collection of MMR signatures defined using samples from individuals with CMMRD (Sanders et al (22)). The Sanders study identified six component signatures (MMR1-6) made up of both substitutions and indels. It included MMR-1, a signature that is predominantly made up of methylation damage and resembles COSMIC signature SBS1. Since these signatures were based on tissues lacking MMR, we focused our analysis on Msh6-deficient samples. *Msh6^−/−^*and *Mbd4^−/−^Msh6^−/−^* bone marrow cells were dominated by MMR-5 and MMR-6 (combined representation of MMR-5/6 was 66% and 67%, respectively) (Figure 4C-D), which are mostly made up of small deletions. There was no significant difference between the proportions or number of mutations attributed to each MMR signature between *Msh6^−/−^* and *Mbd4^−/−^Msh6^−/−^*progenitors, either in bone marrow or colon. The cell lines exhibited an even more marked increase in the absolute number of MMR-5/6 mutations, whereas MMR-3 mutations (replication lagging) were enriched in the thymomas. Despite the differences between tissues, the overall patterns were consistent between *Msh6^−/−^* and *Mbd4^−/−^Msh6^−/−^* samples.

In order to quantify the difference in methylation damage between the various tissues, we studied the rate of MMR1 mutations in primary (Figure 4E) and transformed (Figure 4F) cells of all genotypes. As expected, we consistently saw a marked increase of MMR-1 in *Mbd4^−/−^* samples compared to wild type, which increased further in *Msh6^−/−^* cells. There was no major increase in MMR1 between the *Msh6^−/−^* single and *Mbd4^−/−^ Msh6^−/−^* double knockouts (Figure 4E-F). On the whole, the results largely reflected what we had seen with CG>TG mutations (Figure 2D & 3D), but this approach accounts for overlapping signatures related to MMRd.

Given there was an increase in CG>TG damage in both *Mbd4^−/−^* and *Msh6^−/−^* cells, we sought to determine whether the characteristics and distribution of this damage were consistent. We compared the methylation status of CG>TG mutation sites to simulations of random CG sites (Figure 4G). We found that CG>TG mutations were more likely to occur at methylated sites compared to random, and this was statistically significant in all tissues except for the wild type bone marrow and colon progenitors, which may be due to the low numbers of mutations. To provide more observations for testing, we sourced a set of published somatic mutation calls from colonic crypts (6) from wild type mice, labelled here as “WT crypts”, for use in later comparisons.

We tested for expected differences in replication strand bias between MMR-1-6. In accordance with the human data, T>C mutations, which are mainly found in MMR-3, showed a strong lagging strand bias in all *Msh6^−/−^*tissues (Figure 4H). There were comparatively few T>C mutations in wild type or *Mbd4^−/−^* samples, though they showed no strand bias (Figure 4H). CG>TG mutations in MMR-1 show no replication strand bias in CMMRD samples (22). We found little to no replication strand bias for CG>TG mutations in primary samples, however there was some bias towards the leading strand in the thymomas and cell lines (Figure 4H), which is consistent with a higher contribution from MMR-2-4 and MMR-6.

Replication timing analysis showed the expected enrichment of mutations in late replicating regions (Figure 4I). For methylation damage, the enrichment for late replicating regions was much stronger in the wild type than in *Mbd4^−/−^* or *Msh6^−/−^* tissues. Indeed, loss of Msh6 was associated with less replication timing bias, similar to the observed pattern in human tissues with loss of MSH6 or MSH2.

### Signature development over time

Work with CMMRD tissues has suggested some ability to assess the relative contribution of methylation damage and replicative damage by looking separately at the four NCG>NTG trimer transitions. Looking at the profile of NCG>NTG trimers (Figure 5A), we found that overall, there was good concordance between our mouse models and human tissues. For this comparison we selected *MSH6* mutated cancers from TCGA enriched for methylation damage. Mutations in the GCG>GTG context were much more prevalent in *Msh6^−/−^* cells with high levels of proliferation (Figure 5A, mouse thymoma and MSH6-deficient HAP1 cells), because this trimer is dominant in MMR-2, a replicative signature (22).

**Figure 5:**
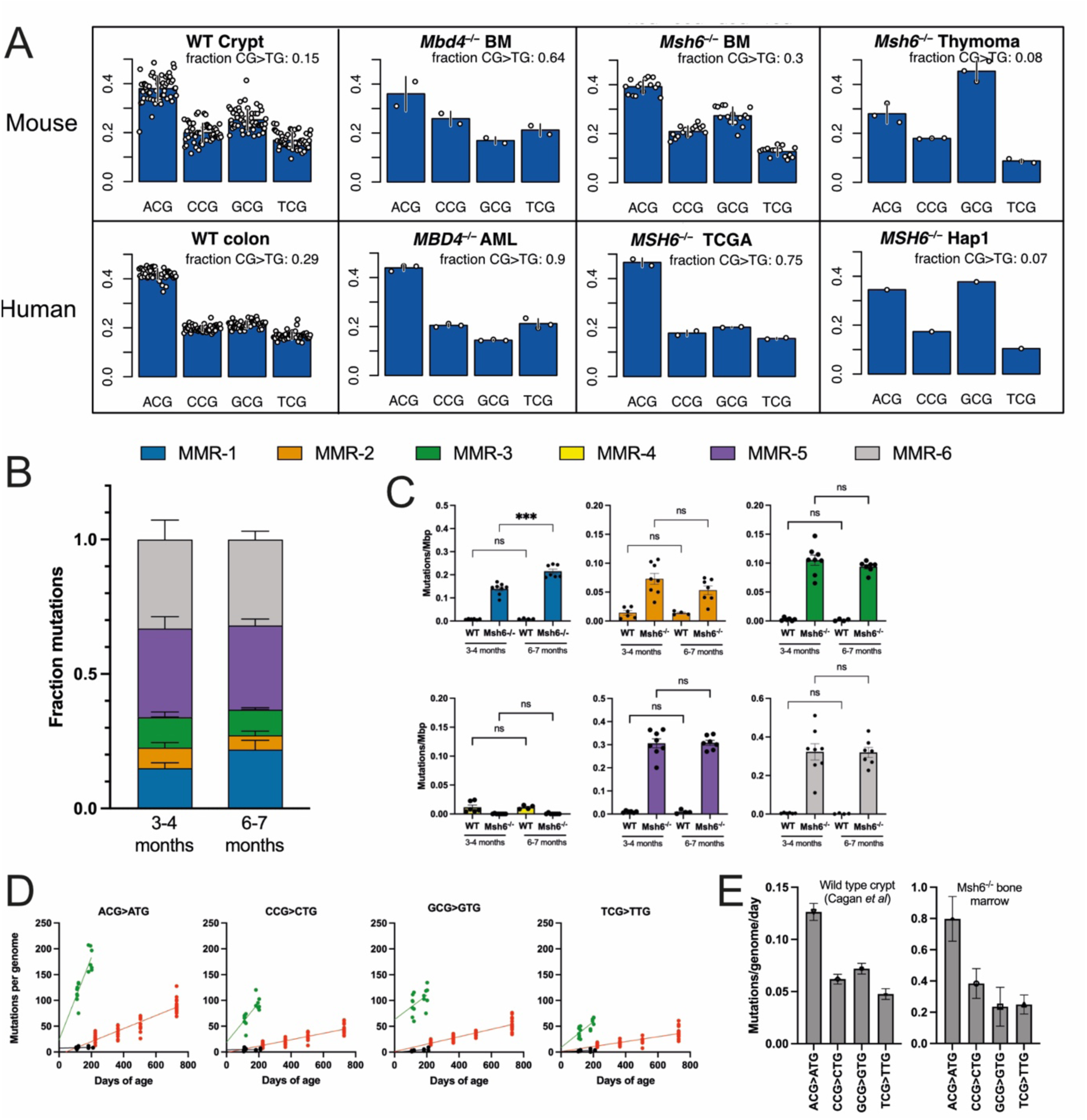
Investigating the impact of cell proliferation and ageing on methylation damage signatures. (A) The proportion of methylation damage spread across each NCG>NTG trimer, for mouse and human samples with different genotypes. Rates are corrected to the trimer frequencies of mouse genomes outside repeat regions. TCGA samples designated MSH6-deficient based on predicted deleterious variants with >80% CG>TG mutations. The mean representation of CG>TG mutations is calculated for sample group. Each datapoint represents a sample and error bars represent SD. (B) MMR signature deconvolution from somatic mutations from 3-4 month and 6-7 month bone marrow progenitors. Data for 3-4 month old samples is the same as plotted in Figure 4D and E. (C) Rates of each of the 6 MMR signatures, split by genotype and by age. Each datapoint represents a colony with mean and SD shown. Significance tested in Prism with ANOVA with Welch’s correction ( *** p<0.001) (D) Accumulation of methylation damage over time, considering each NCG>NTG trimer, in 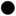 wild type bone marrow, 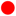 wild type crypt (6), or 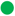 *Msh6^−/−^* bone marrow. Each point represents a colony or sample. A linear regression line is fit. (E) Accumulation of mutations in NCG context over time, plotted as mutations/genome/day. Error represents standard error of the slope of the respective linear regressions in D.

We sought to determine whether methylation damage would accumulate independent of the replicative damage in our models. To do this we compared mutational signatures in bone marrow progenitors from *Msh6^−/−^* mice aged for 6-7 months (n=7) to our existing dataset from 3 month old mice (n=8). We found that the fraction of MMR-1 increased in the older mice (Figure 5B). Looking at mutation rate of each MMR signature (Figure 5C), we found that MMR-1 increased significantly (from 0.14 to 0.21 mutations/Mbp, p =0.0002, one way ANOVA with Welches correction), but all other MMR signatures remained stable, with no significant increases. This finding is also evident when looking at CG>TG mutations, compared to other SBS or indels (Figure S7). These findings are consistent with an early burst of replicative damage, and steady accumulation of methylation damage. We decided to assess the linear fit for mutation rate for each NCG>NTG transition in wild type and Msh6-deficient samples (Figure 5D). Despite the differing slopes on the linear regressions, all of the wild type slopes intersect the x-axis around the point of conception (e0 or 19-21 days before birth), suggesting a consistent rate across the lifetime. This is similar for three of the trimers in *Msh6^−/−^* bone marrow, albeit with a steeper slope. The GCG>GTG mutations were the exception; the non-zero intersection at the time of conception indicates a surge of mutations before the 3-month timepoint. This is consistent with a burst of replicative damage during development. GCG>GTG mutations then increase at a rate similar to CCG>CTG and TCG>TTG suggesting that after 3 months of age, GCG>GTG mutations accumulate similarly to other types of methylation damage (Figure 5E).

## Discussion

The chemical properties of 5mC make it prone to deamination (48). As a result, DNA repair mechanisms have developed alongside 5mC, to counter its mutagenic influence. This form of DNA repair is somewhat specialised, in that, it should always be directed towards removal of the thymine from the G/T mismatch and, since thymine is a normal component of DNA, its removal needs to be strictly controlled. Our work has highlighted a key role for the DNA glycosylase MBD4 in mediating the repair of methylation damage in humans and mice (7,9). The use of a DNA glycosylase provides specificity and orients the repair activity to remove the thymine, but this is not the only way to approach this problem. In *Escherichia coli* strains that use 5mC, the repair of methylation damage is initiated by a sequence and methylation state specific endonuclease, Vsr (49,50). Vsr orients the repair towards the thymine, in a process termed very short patch repair (VSP), which also relies on PolA, MutS and MutL (reviewed in (51)). It is notable that three recent genomics studies have implicated MMR components in the repair of methylation damage in people (20,22,52). We set out to determine which DNA glycosylases contribute to the repair of methylation damage in the mammalian genome, whether they interact with the MMR pathway and whether there are direct parallels with DNA repair strategies in other species.

TDG was the first glycosylase implicated in methylation damage repair; it was identified based on its ability to bind a G/T mismatch and excise thymine (15,40,41). Yet, initial studies with Tdg knockout mice failed to reveal elevated rates of somatic mutation, and instead identified a role in modulating DNA methylation and transcription (17,53). It was subsequently revealed that TDG contributes to active demethylation through excision of 5-formyl- and 5-carboxy-cytosine, the oxidative products of 5mC (54–56). More recent profiling of genome-wide occupancy and protein-protein interaction studies also position TDG as a transcriptional regulator (57). Nonetheless, given the overlapping substrate preference, we sought to determine whether Tdg functionally compensates for the loss of Mbd4. However, we found no evidence of elevated methylation damage in Tdg-deficient cells. Combined loss of Tdg and Mbd4 resulted in no significant increase beyond the levels of methylation damage seen with Mbd4 loss alone. While the level of methylation damage was slightly higher in compound mutant cell lines, this was not significant and occurred in the background of a higher overall mutation rate, which could reflect differences between clones. Our findings align with results obtained with genome sequencing performed in human TDG-deficient induced pluripotent stem cells (18), and extend them in two meaningful ways. First, our experimental design was more suited to detect methylation damage, as it leveraged primary cells and looked over longer time periods. Second, by assessing double mutant cells, we could confirm that Tdg does not compensate for the loss of Mbd4. These results align with a recently published effort, which look at the impact of combined loss of TDG and MBD4 in HAP1 cell lines (58). As there is ample biochemical evidence that Tdg can excise thymine, these results suggest its enzymatic activity is restricted within cells, which may reflect the influence of regulatory mechanisms, such as post-translational modifications (59), or that it is not being recruited effectively to sites of methylation damage.

Several recent studies have implicated MMR, and specifically the MutSα complex (MSH6:MSH2) in methylation damage repair in humans (20,22,52). These studies have used signature deconvolution to help separate classic replication-coupled MMR from repair activity outside replication, sometimes referred to as non-canonical MMR. We began by examining the mutational signature of *Msh6^−/−^*progenitor cells from the bone marrow and colon. Many features of the MMRd signature were consistent between mice and humans, but the substitution to indel ratio was a point of difference. The mutational signatures from humans showed higher numbers of substitutions relative to indels, and some showed marked accumulation of methylation damage (≥60% of all SBS), a phenomenon referred to as somatic CG hypermutation (20,22,52). Given that methylation damage results from spontaneous deamination, it is not surprising to see stronger enrichment in people, as there is more time for the damage to accumulate compared to laboratory models. By aging mice further, we found that methylation damage continued to accumulate in bone marrow progenitors between 3 and 6 months, whereas the amount of replicative damage seemed stable. This would fit a model in which the replicative damage in primary bone marrow cells accumulated early, whereas the methylation damage continued to accumulate at a relatively constant rate, even in quiescent cells. It also highlights the challenge of observing an enrichment in methylation damage in rapidly dividing cells, where the vast majority of damage will be due to replicative errors. 5mC can pose challenges for the replication machinery, for example, a recent assessment of DNA polymerases showed that CG methylation resulted in a seven-fold increase in C>T errors (60). Indeed, replicative signatures became dominant in our Msh6-deficient cell lines and thymomas, which mirrors prior work in cell lines (18). It is notable that CG hypermutation is not evident in MMR-deficient yeast (61) or worms (62), which lack endogenous 5mC. Collectively, these studies support the view that the replication-coupled and non-canonical DNA repair activities can be separated, and that mutational signatures detected in MMRd human tissues strongly reflect the replicative history of the cell.

In classic MMR, MutSα recruits MutL to initiate endonucleolytic cleavage and instigate strand resynthesis. This repair activity is tightly regulated during the cell cycle (63) and it relies on differences in methylation status to help orient repair activity – these features are ill suited to repair spontaneous methylation damage. Furthermore, even though MSH6 and MBD4 can both interact with MLH1, the evidence from human cancers and CMMRD suggests that MutL is not involved in methylation damage repair (20,22). We propose that instead, MutSα and MBD4 cooperate directly in the identification and repair of methylation damage. Mismatch proteins including Msh6 have been shown to be expressed throughout G_0_/G_1_ as well as S phase (64,65), and it is notable that both MBD4 and MSH6 can bind G/T mismatches *in vitro* (21,66). By recruiting MBD4, the resulting MutSα-MBD4 complex would gain selectivity for removal of thymine, analogous to the VSP repair process in *E.coli*. Furthermore, it has been shown that the thymine glycosylase activity of MBD4 is enhanced through interaction with MMR components, and it is further stimulated by inclusion of 5mC in the template DNA, which could help orient repair (67). This suggestion of cooperation is supported by the observation that we did not see a marked increase in the levels of methylation damage when we combined loss of Mbd4 and Msh6, beyond that seen with Msh6-deficiency alone. Cells that lack Mbd4 acquire about half as much methylation damage as Msh6-deficient cells, suggesting that MutSα may have some ability to identify methylation damage and initiate repair independent of Mbd4, but that repair may be less efficient, or it may be oriented to the wrong strand some of the time. If MMR components recruit or activate MBD4, it may explain why Tdg, despite having thymine glycosylase activity *in vitro*, does not have this activity *in vivo*. These additional licensing steps may act as a safeguard, helping to prevent unnecessary or erroneous excision of thymine.

Prior work on CG hypermutation in human cancers has suggested a potential role for TDG in guiding non-canonical MMR. Our results instead point to cooperation between MutSα and Mbd4. There are limitations to our study. First, we assume that excessive methylation damage in the context of MMRd reflects the situation in which cells experience an extended period of dormancy, where methylation damage accumulates without significant replicative damage, but we were only able to model this process over 3-6 months. It would be ideal to perform longer ageing experiments and to extend these studies to look at other tissues, including post-replicative tissues. The targeted allele could be used to induce *Msh6* deletion in adult mice, to substantiate the hypothesis of a strong burst of replicative damage in early development. Mutational profiling could also be pursued using error corrected sequencing which would allow investigation of many tissues without the need for clonal expansion (68). Our knockout allele results in complete loss of the Msh6 protein and destabilisation of Msh2. Human CMMRD patients may have complete loss, or they may carry severe loss of function mutations, where they retain some stable protein (69). This might contribute to some of the variation in mutation profiles seen in individuals with germline or somatic loss of MSH6, or it could also reflect the timing of MSH6 loss relative to transformation. Finally, we provide genetic evidence for interaction between MutSα and MBD4, but this should be investigated using biochemical and structural approaches.

Our results position MBD4 and MutSα as the main safeguards against methylation damage for the mammalian genome. As methylation damage is a major contributor to both somatic and germline mutagenesis, strategies to enhance the functional activity of these complexes may offer a way to slow the degradation of the genome, providing for a longer, healthier life.

## Supporting information

Supplementary Figures 1-7 and description of Supplementary Tables.

## Data availability

Descriptive statistics regarding coverage assessments and mutation rates are provided in the supplement (Table S4-5). Analytical code to reproduce figure elements has been deposited through Figshare, along with a set of processed variant calls and a mutation matrix (DOI: 10.6084/m9.figshare.27998555). Aligned sequencing data will be made available for research use through the Sequence Read Archive.

## Acknowledgements

We thank Maree Faux, Sandra Mifsud and Ladina Di Rago for technical assistance, Artem Laktyushin and Jeff Babon for producing TDG protein and the WEHI Antibody Facility for development of monoclonal antibodies. We wish to acknowledge the WEHI Bioservices team for care of experimental mice and the WEHI Histology team for assistance processing tissue samples. We thank Jason Wong for valuable feedback. The authors acknowledge the facilities, and the scientific and technical assistance of The Melbourne Advanced Genome Editing Centre (MAGEC). DNA sequencing services were provided by the Australian Genome Research Facility (AGRF) and BGI Genomics. Analysis of human genomic data was approved by the WEHI Human Research Ethics Committee (Project 13-01 IJM). The results are based, in part, on data generated by the TCGA Research Network (http://cancergenome.nih.gov/).

## References

1. Chatterjee, N. and Walker, G.C. (2017) Mechanisms of DNA damage, repair, and mutagenesis. Environ Mol Mutagen, 58, 235–263.

2. Blokzijl, F., de Ligt, J., Jager, M., Sasselli, V., Roerink, S., Sasaki, N., Huch, M., Boymans, S., Kuijk, E., Prins, P., et al. (2016) Tissue-specific mutation accumulation in human adult stem cells during life. Nature, 538, 260–264.

3. Alexandrov, L.B., Nik-Zainal, S., Wedge, D.C., Aparicio, S.A., Behjati, S., Biankin, A.V., Bignell, G.R., Bolli, N., Borg, A., Borresen-Dale, A.L. et al. (2013) Signatures of mutational processes in human cancer. Nature, 500, 415–421.

4. Rahbari, R., Wuster, A., Lindsay, S.J., Hardwick, R.J., Alexandrov, L.B., Al Turki, S., Dominiczak, A., Morris, A., Porteous, D., Smith, B., et al. (2016) Timing, rates and spectra of human germline mutation. Nat Genet, 48, 126–133.

5. Alexandrov, L.B., Kim, J., Haradhvala, N.J., Huang, M.N., Tian Ng, A.W., Wu, Y., Boot, A., Covington, K.R., Gordenin, D.A., Bergstrom, E.N. et al. (2020) The repertoire of mutational signatures in human cancer. Nature, 578, 94–101.

6. Cagan, A., Baez-Ortega, A., Brzozowska, N., Abascal, F., Coorens, T.H.H., Sanders, M.A., Lawson, A.R.J., Harvey, L.M.R., Bhosle, S., Jones, D. et al. (2022) Somatic mutation rates scale with lifespan across mammals. Nature, 604, 517–524.

7. Sanders, M.A., Chew, E., Flensburg, C., Zeilemaker, A., Miller, S.E., Al Hinai, A.S., Bajel, A., Luiken, B., Rijken, M., McLennan, T., et al. (2018) MBD4 guards against methylation damage and germ line deficiency predisposes to clonal hematopoiesis and early-onset AML. Blood, 132, 1526–1534.

8. Derrien, A.C., Rodrigues, M., Eeckhoutte, A., Dayot, S., Houy, A., Mobuchon, L., Gardrat, S., Lequin, D., Ballet, S., Pierron, G. et al. (2021) Germline MBD4 Mutations and Predisposition to Uveal Melanoma. J Natl Cancer Inst, 113, 80–87.

9. Palles, C., West, H.D., Chew, E., Galavotti, S., Flensburg, C., Grolleman, J.E., Jansen, E.A.M., Curley, H., Chegwidden, L., Arbe-Barnes, E.H. et al. (2022) Germline MBD4 deficiency causes a multi-tumor predisposition syndrome. Am J Hum Genet, 109, 953–960.

10. Rodrigues, M., Mobuchon, L., Houy, A., Fievet, A., Gardrat, S., Barnhill, R.L., Popova, T., Servois, V., Rampanou, A., Mouton, A. et al. (2018) Outlier response to anti-PD1 in uveal melanoma reveals germline MBD4 mutations in hypermutated tumors. Nat Commun, 9, 1866.

11. Hendrich, B., Hardeland, U., Ng, H.H., Jiricny, J. and Bird, A. (1999) The thymine glycosylase MBD4 can bind to the product of deamination at methylated CpG sites. Nature, 401, 301–304.

12. Millar, C.B., Guy, J., Sansom, O.J., Selfridge, J., MacDougall, E., Hendrich, B., Keightley, P.D., Bishop, S.M., Clarke, A.R. and Bird, A. (2002) Enhanced CpG mutability and tumorigenesis in MBD4-deficient mice. Science, 297, 403–405.

13. Wong, E., Yang, K., Kuraguchi, M., Werling, U., Avdievich, E., Fan, K., Fazzari, M., Jin, B., Brown, A.M., Lipkin, M. et al. (2002) Mbd4 inactivation increases C>T transition mutations and promotes gastrointestinal tumor formation. Proc Natl Acad Sci U S A, 99, 14937–14942.

14. Shen, J.C., Rideout, W.M., 3rd and Jones, P.A. (1994) The rate of hydrolytic deamination of 5-methylcytosine in double-stranded DNA. Nucleic Acids Res, 22, 972–976.

15. Yoon, J.H., Iwai, S., O’Connor, T.R. and Pfeifer, G.P. (2003) Human thymine DNA glycosylase (TDG) and methyl-CpG-binding protein 4 (MBD4) excise thymine glycol (Tg) from a Tg:G mispair. Nucleic Acids Res, 31, 5399–5404.

16. Maiti, A. and Drohat, A.C. (2011) Thymine DNA glycosylase can rapidly excise 5-formylcytosine and 5-carboxylcytosine: potential implications for active demethylation of CpG sites. J Biol Chem, 286, 35334–35338.

17. Cortazar, D., Kunz, C., Selfridge, J., Lettieri, T., Saito, Y., MacDougall, E., Wirz, A., Schuermann, D., Jacobs, A.L., Siegrist, F. et al. (2011) Embryonic lethal phenotype reveals a function of TDG in maintaining epigenetic stability. Nature, 470, 419–423.

18. Zou, X., Koh, G.C.C., Nanda, A.S., Degasperi, A., Urgo, K., Roumeliotis, T.I., Agu, C.A., Badja, C., Momen, S., Young, J. et al. (2021) A systematic CRISPR screen defines mutational mechanisms underpinning signatures caused by replication errors and endogenous DNA damage. Nat Cancer, 2, 643–657.

19. Kunkel, T.A. and Erie, D.A. (2015) Eukaryotic Mismatch Repair in Relation to DNA Replication. Annu Rev Genet, 49, 291–313.

20. Fang, H., Zhu, X., Yang, H., Oh, J., Barbour, J.A. and Wong, J.W.H. (2021) Deficiency of replication-independent DNA mismatch repair drives a 5-methylcytosine deamination mutational signature in cancer. Sci Adv, 7, eabg4398.

21. Palombo, F., Gallinari, P., Iaccarino, I., Lettieri, T., Hughes, M., D’Arrigo, A., Truong, O., Hsuan, J.J. and Jiricny, J. (1995) GTBP, a 160-kilodalton protein essential for mismatch-binding activity in human cells. Science, 268, 1912–1914.

22. Sanders, M.A., Vöhringer, H., Forster, V.J., Moore, L., Campbell, B.B., Hooks, Y., Edwards, M., Bianchi, V., Coorens, T.H.H., Butler, T.M., et al. (2021) Life without mismatch repair. BioRxiv, https://www.biorxiv.org/content/10.1101/2021.1104.1114.437578v437571.

23. de Boer, J., Williams, A., Skavdis, G., Harker, N., Coles, M., Tolaini, M., Norton, T., Williams, K., Roderick, K., Potocnik, A.J., et al. (2003) Transgenic mice with hematopoietic and lymphoid specific expression of Cre. Eur J Immunol, 33, 314–325.

24. Ventura, A., Kirsch, D.G., McLaughlin, M.E., Tuveson, D.A., Grimm, J., Lintault, L., Newman, J., Reczek, E.E., Weissleder, R. and Jacks, T. (2007) Restoration of p53 function leads to tumour regression in vivo. Nature, 445, 661–665.

25. Hirokawa, Y., Yip, K.H., Tan, C.W. and Burgess, A.W. (2014) Colonic myofibroblast cell line stimulates colonoid formation. Am J Physiol Gastrointest Liver Physiol, 306, G547–556.

26. Yip, H.Y.K., Tan, C.W., Hirokawa, Y. and Burgess, A.W. (2018) Colon organoid formation and cryptogenesis are stimulated by growth factors secreted from myofibroblasts. PLoS One, 13, e0199412.

27. Xu, Z., Flensburg, C., Bilardi, R.A. and Majewski, I.J. (2023) Uridine-cytidine kinase 2 potentiates the mutagenic influence of the antiviral beta-d-N4-hydroxycytidine. Nucleic Acids Res, 51, 12031–12042.

28. Aubrey, B.J., Kelly, G.L., Kueh, A.J., Brennan, M.S., O’Connor, L., Milla, L., Wilcox, S., Tai, L., Strasser, A. and Herold, M.J. (2015) An inducible lentiviral guide RNA platform enables the identification of tumor-essential genes and tumor-promoting mutations in vivo. Cell Rep, 10, 1422–1432.

29. Li, H. and Durbin, R. (2009) Fast and accurate short read alignment with Burrows-Wheeler transform. Bioinformatics, 25, 1754–1760.

30. (2019) Picard toolkit. Broad Institute, GitHub repository.

31. Flensburg, C., Sargeant, T., Oshlack, A. and Majewski, I.J. (2020) SuperFreq: Integrated mutation detection and clonal tracking in cancer. PLoS Comput Biol, 16, e1007603.

32. Smit, A., Hubley, R & Green, P. (1996-2010) RepeatMasker Open-3.0. http://www.repeatmasker.org.

33. Obenchain, V., Lawrence, M., Carey, V., Gogarten, S., Shannon, P. and Morgan, M. (2014) VariantAnnotation: a Bioconductor package for exploration and annotation of genetic variants. Bioinformatics, 30, 2076–2078.

34. Tate, J.G., Bamford, S., Jubb, H.C., Sondka, Z., Beare, D.M., Bindal, N., Boutselakis, H., Cole, C.G., Creatore, C., Dawson, E. et al. (2019) COSMIC: the Catalogue Of Somatic Mutations In Cancer. Nucleic Acids Res, 47, D941–D947.

35. Manders, F., Brandsma, A.M., de Kanter, J., Verheul, M., Oka, R., van Roosmalen, M.J., van der Roest, B., van Hoeck, A., Cuppen, E. and van Boxtel, R. (2022) MutationalPatterns: the one stop shop for the analysis of mutational processes. BMC Genomics, 23, 134.

36. Tubbs, A., Sridharan, S., van Wietmarschen, N., Maman, Y., Callen, E., Stanlie, A., Wu, W., Wu, X., Day, A., Wong, N., et al. (2018) Dual Roles of Poly(dA:dT) Tracts in Replication Initiation and Fork Collapse. Cell, 174, 1127–1142 e1119.

37. Ryba, T., Battaglia, D., Pope, B.D., Hiratani, I. and Gilbert, D.M. (2011) Genome-scale analysis of replication timing: from bench to bioinformatics. Nat Protoc, 6, 870–895.

38. Hon, G.C., Rajagopal, N., Shen, Y., McCleary, D.F., Yue, F., Dang, M.D. and Ren, B. (2013) Epigenetic memory at embryonic enhancers identified in DNA methylation maps from adult mouse tissues. Nat Genet, 45, 1198–1206.

39. Moore, L., Cagan, A., Coorens, T.H.H., Neville, M.D.C., Sanghvi, R., Sanders, M.A., Oliver, T.R.W., Leongamornlert, D., Ellis, P., Noorani, A. et al. (2021) The mutational landscape of human somatic and germline cells. Nature, 597, 381–386.

40. Neddermann, P. and Jiricny, J. (1993) The purification of a mismatch-specific thymine-DNA glycosylase from HeLa cells. J Biol Chem, 268, 21218–21224.

41. Neddermann, P., Gallinari, P., Lettieri, T., Schmid, D., Truong, O., Hsuan, J.J., Wiebauer, K. and Jiricny, J. (1996) Cloning and expression of human G/T mismatch-specific thymine-DNA glycosylase. J Biol Chem, 271, 12767–12774.

42. Onodera, A., Gonzalez-Avalos, E., Lio, C.J., Georges, R.O., Bellacosa, A., Nakayama, T. and Rao, A. (2021) Roles of TET and TDG in DNA demethylation in proliferating and non-proliferating immune cells. Genome Biol, 22, 186.

43. Goncalves, E., Poulos, R.C., Cai, Z., Barthorpe, S., Manda, S.S., Lucas, N., Beck, A., Bucio-Noble, D., Dausmann, M., Hall, C. et al. (2022) Pan-cancer proteomic map of 949 human cell lines. Cancer Cell, 40, 835–849 e838.

44. Bellacosa, A., Cicchillitti, L., Schepis, F., Riccio, A., Yeung, A.T., Matsumoto, Y., Golemis, E.A., Genuardi, M. and Neri, G. (1999) MED1, a novel human methyl-CpG-binding endonuclease, interacts with DNA mismatch repair protein MLH1. Proc Natl Acad Sci U S A, 96, 3969–3974.

45. Cortellino, S., Turner, D., Masciullo, V., Schepis, F., Albino, D., Daniel, R., Skalka, A.M., Meropol, N.J., Alberti, C., Larue, L. et al. (2003) The base excision repair enzyme MED1 mediates DNA damage response to antitumor drugs and is associated with mismatch repair system integrity. Proc Natl Acad Sci U S A, 100, 15071–15076.

46. Edelmann, W., Yang, K., Umar, A., Heyer, J., Lau, K., Fan, K., Liedtke, W., Cohen, P.E., Kane, M.F., Lipford, J.R. et al. (1997) Mutation in the mismatch repair gene Msh6 causes cancer susceptibility. Cell, 91, 467–477.

47. Sansom, O.J., Bishop, S.M., Bird, A. and Clarke, A.R. (2004) MBD4 deficiency does not increase mutation or accelerate tumorigenesis in mice lacking MMR. Oncogene, 23, 5693–5696.

48. Zhang, X. and Mathews, C.K. (1994) Effect of DNA cytosine methylation upon deamination-induced mutagenesis in a natural target sequence in duplex DNA. J Biol Chem, 269, 7066–7069.

49. Sohail, A., Lieb, M., Dar, M. and Bhagwat, A.S. (1990) A gene required for very short patch repair in Escherichia coli is adjacent to the DNA cytosine methylase gene. J Bacteriol, 172, 4214–4221.

50. Hennecke, F., Kolmar, H., Brundl, K. and Fritz, H.J. (1991) The vsr gene product of E. coli K-12 is a strand- and sequence-specific DNA mismatch endonuclease. Nature, 353, 776–778.

51. Bhagwat, A.S. and Lieb, M. (2002) Cooperation and competition in mismatch repair: very short-patch repair and methyl-directed mismatch repair in Escherichia coli. Mol Microbiol, 44, 1421–1428.

52. Flynn, A., Waszak, S.M. and Weischenfeldt, J. (2024) Somatic CpG hypermutation is associated with mismatch repair deficiency in cancer. Mol Syst Biol.

53. Cortellino, S., Xu, J., Sannai, M., Moore, R., Caretti, E., Cigliano, A., Le Coz, M., Devarajan, K., Wessels, A., Soprano, D., et al. (2011) Thymine DNA glycosylase is essential for active DNA demethylation by linked deamination-base excision repair. Cell, 146, 67–79.

54. He, Y.F., Li, B.Z., Li, Z., Liu, P., Wang, Y., Tang, Q., Ding, J., Jia, Y., Chen, Z., Li, L. et al. (2011) Tet-mediated formation of 5-carboxylcytosine and its excision by TDG in mammalian DNA. Science, 333, 1303–1307.

55. Ito, S., Shen, L., Dai, Q., Wu, S.C., Collins, L.B., Swenberg, J.A., He, C. and Zhang, Y. (2011) Tet proteins can convert 5-methylcytosine to 5-formylcytosine and 5-carboxylcytosine. Science, 333, 1300–1303.

56. Raiber, E.A., Beraldi, D., Ficz, G., Burgess, H.E., Branco, M.R., Murat, P., Oxley, D., Booth, M.J., Reik, W. and Balasubramanian, S. (2012) Genome-wide distribution of 5-formylcytosine in embryonic stem cells is associated with transcription and depends on thymine DNA glycosylase. Genome Biol, 13, R69.

57. Aranda, S., Alcaine-Colet, A., Ballare, C., Blanco, E., Mocavini, I., Sparavier, A., Vizan, P., Borras, E., Sabido, E. and Di Croce, L. (2023) Thymine DNA glycosylase regulates cell-cycle-driven p53 transcriptional control in pluripotent cells. Mol Cell, 83, 2673–2691 e2677.

58. Silveira, A.B., Houy, A., Ganier, O., Ozemek, B., Vanhuele, S., Vincent-Salomon, A., Cassoux, N., Mariani, P., Pierron, G., Leyvraz, S. et al. (2024) Base-excision repair pathway shapes 5-methylcytosine deamination signatures in pan-cancer genomes. Nat Commun, 15, 9864.

59. Pidugu, L.S., Servius, H.W., Espinosa, K.B., Cook, M.E., Varney, K.M. and Drohat, A.C. (2024) Sumoylation of thymine DNA glycosylase impairs productive binding to substrate sites in DNA. J Biol Chem, 300, 107902.

60. Tomkova, M., McClellan, M.J., Crevel, G., Shahid, A.M., Mozumdar, N., Tomek, J., Shepherd, E., Cotterill, S., Schuster-Bockler, B. and Kriaucionis, S. (2024) Human DNA polymerase epsilon is a source of C>T mutations at CpG dinucleotides. Nat Genet.

61. Kadyrova, L.Y., Mieczkowski, P.A. and Kadyrov, F.A. (2023) Genome-wide contributions of the MutSalpha- and MutSbeta-dependent DNA mismatch repair pathways to the maintenance of genetic stability in Saccharomyces cerevisiae. J Biol Chem, 299, 104705.

62. Meier, B., Volkova, N.V., Hong, Y., Schofield, P., Campbell, P.J., Gerstung, M. and Gartner, A. (2018) Mutational signatures of DNA mismatch repair deficiency in C. elegans and human cancers. Genome Res, 28, 666–675.

63. Rona, G., Miwatani-Minter, B., Zhang, Q., Goldberg, H.V., Kerzhnerman, M.A., Howard, J.B., Simoneschi, D., Lane, E., Hobbs, J.W., Sassani, E. et al. (2024) CDK-independent role of D-type cyclins in regulating DNA mismatch repair. Mol Cell, 84, 1224–1242 e1213.

64. Edelbrock, M.A., Kaliyaperumal, S. and Williams, K.J. (2009) DNA mismatch repair efficiency and fidelity are elevated during DNA synthesis in human cells. Mutat Res, 662, 59–66.

65. Schroering, A.G., Edelbrock, M.A., Richards, T.J. and Williams, K.J. (2007) The cell cycle and DNA mismatch repair. Exp Cell Res, 313, 292–304.

66. Petronzelli, F., Riccio, A., Markham, G.D., Seeholzer, S.H., Genuardi, M., Karbowski, M., Yeung, A.T., Matsumoto, Y. and Bellacosa, A. (2000) Investigation of the substrate spectrum of the human mismatch-specific DNA N-glycosylase MED1 (MBD4): fundamental role of the catalytic domain. J Cell Physiol, 185, 473–480.

67. Papin, C., Ibrahim, A., Sabir, J.S.M., Le Gras, S., Stoll, I., Albiheyri, R.S., Zari, A.T., Bahieldin, A., Bellacosa, A., Bronner, C., et al. (2023) MBD4 loss results in global reactivation of promoters and retroelements with low methylated CpG density. J Exp Clin Cancer Res, 42, 301.

68. Abascal, F., Harvey, L.M.R., Mitchell, E., Lawson, A.R.J., Lensing, S.V., Ellis, P., Russell, A.J.C., Alcantara, R.E., Baez-Ortega, A., Wang, Y. et al. (2021) Somatic mutation landscapes at single-molecule resolution. Nature, 593, 405–410.

69. Aronson, M., Colas, C., Shuen, A., Hampel, H., Foulkes, W.D., Baris Feldman, H., Goldberg, Y., Muleris, M., Wolfe Schneider, K., McGee, R.B. et al. (2022) Diagnostic criteria for constitutional mismatch repair deficiency (CMMRD): recommendations from the international consensus working group. J Med Genet, 59, 318–327.

