## Supplementary Figures 1-7 and description of Supplementary Tables. for "Mbd4 and MutSα protect cells from spontaneous deamination of 5-methylcytosine"

### Supplementary Material

**Manuscript Title:**
Mbd4 and MutS$\alpha$ protect cells from spontaneous deamination of 5-methylcytosine.

**Authors:**
Rebecca A. Bilardi[1,2], Christoffer Flensburg[1], Zhen Xu[1,2], Emily B. Derrick[3,4], Andrew Kueh[1,5,6] and Ian J Majewski*[1,2].

**Affiliation:**
[1]Blood Cells and Blood Cancer Division, The Walter and Eliza Hall Institute, Parkville 3052, Australia
[2]Department of Medical Biology, University of Melbourne, Parkville, 3052, Australia
[3]Cancer Immunology Program, Peter MacCallum Cancer Centre, Melbourne, 3052, Victoria, Australia
[4]Sir Peter MacCallum Department of Oncology, The University of Melbourne, Parkville, 3052, Victoria, Australia
[5]Olivia Newton-John Cancer Research Institute, Heidelberg, 3084, Australia
[6]School of Cancer Medicine, La Trobe University, Bundoora, 3083, Victoria, Australia

**Description:**
This data supplement includes supplementary figures and a description of supplementary tables. In total there are 7 supplementary figures and 5 supplementary tables. Variant call sets are provided through Figshare (DOI: 10.6084/m9.figshare.27998555).

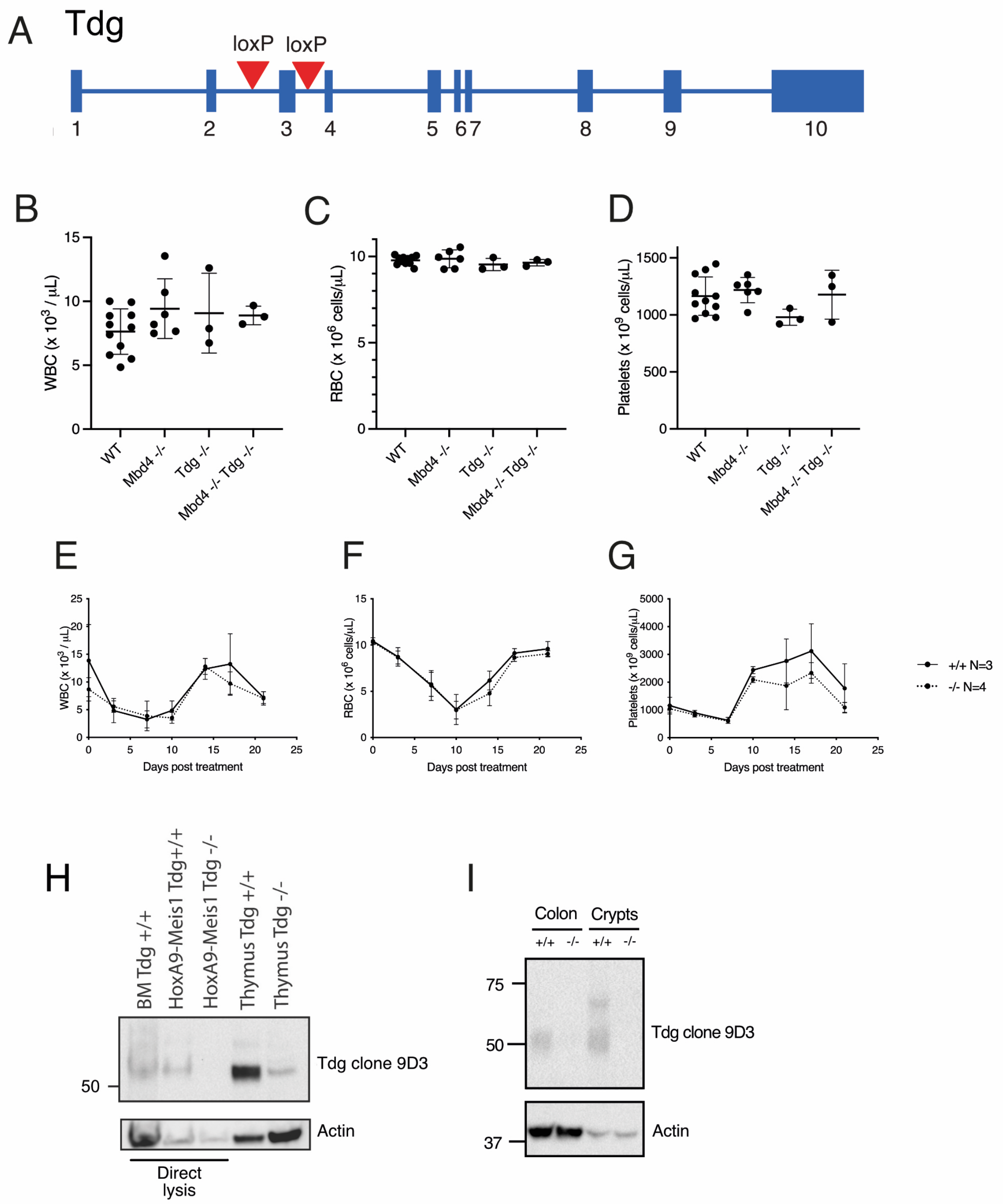


**Supplementary Figure 1: Characterisation of the *Tdg* floxed mouse model.** (A) Schematic of the design of the *Tdg* floxed allele. Deletion of exon 3 results in a premature stop codon and is predicted to give no functional protein. Blood counts at 7 weeks of age via an automated haematological analyser, including (B) white blood cells (WBC), (C) red blood cells (RBC) and (D) platelets, with each showing mean and SD. $Tdg^{fl/+}$ $Vav\text{-}Cre^{+/+}$ (labelled $Tdg^{+/+}$) and $Tdg^{fl/-}$ $Vav\text{-}Cre^{T/+}$ (labelled $Tdg^{-/-}$) mice were treated once with 150 mg/kg 5-fluorouracil intravenously and their blood counts monitored over 3 weeks. Data are presented for (E) WBC, (F) RBC and (G) platelets. (H) Western blotting for Tdg protein. Whole mouse bone marrow or foetal liver derived cell lines were lysed directly in hot SDS PAGE loading buffer. Thymus was lysed in RIPA buffer and quantified prior to electrophoresis. Actin is used as a loading control. Molecular weight is indicated at left (kDa). (I) Western blotting was performed on lysates prepared from whole colon tissue and isolated crypts from tamoxifen treated mice of the genotypes *Tdg*$^{+/+}$ *Rosa26-CreERT2*$^{T/+}$ (labelled *Tdg*$^{+/+}$) or *Tdg*$^{fl/fl}$ *Rosa26-CreERT2*$^{T/+}$ (labelled *Tdg*$^{-/-}$). Actin is used as a loading control. Molecular weight is indicated at left (kDa).

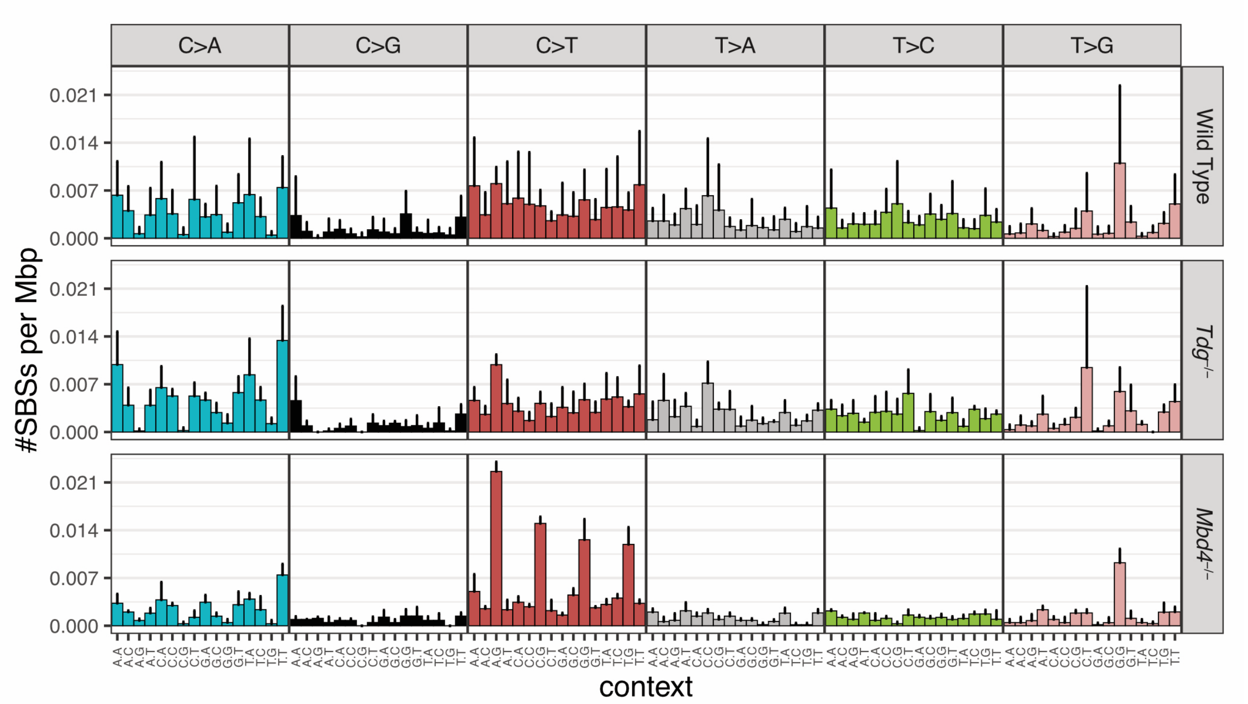


**Supplementary Figure 2: Mutation signature analysis in *Tdg*$^{-/-}$ and *Mbd4*$^{-/-}$ mice at 3 months.** Average mutation signatures of colon derived colonies from wild type (N = 9 colonies), *Rosa-CreERT2*$^{T/+}$ *Tdg*$^{fl/fl}$ (*Tdg*$^{-/-}$) (N = 4 colonies) and *Mbd4*$^{-/-}$ (N = 3 colonies) mice. All values represent mean and SD.

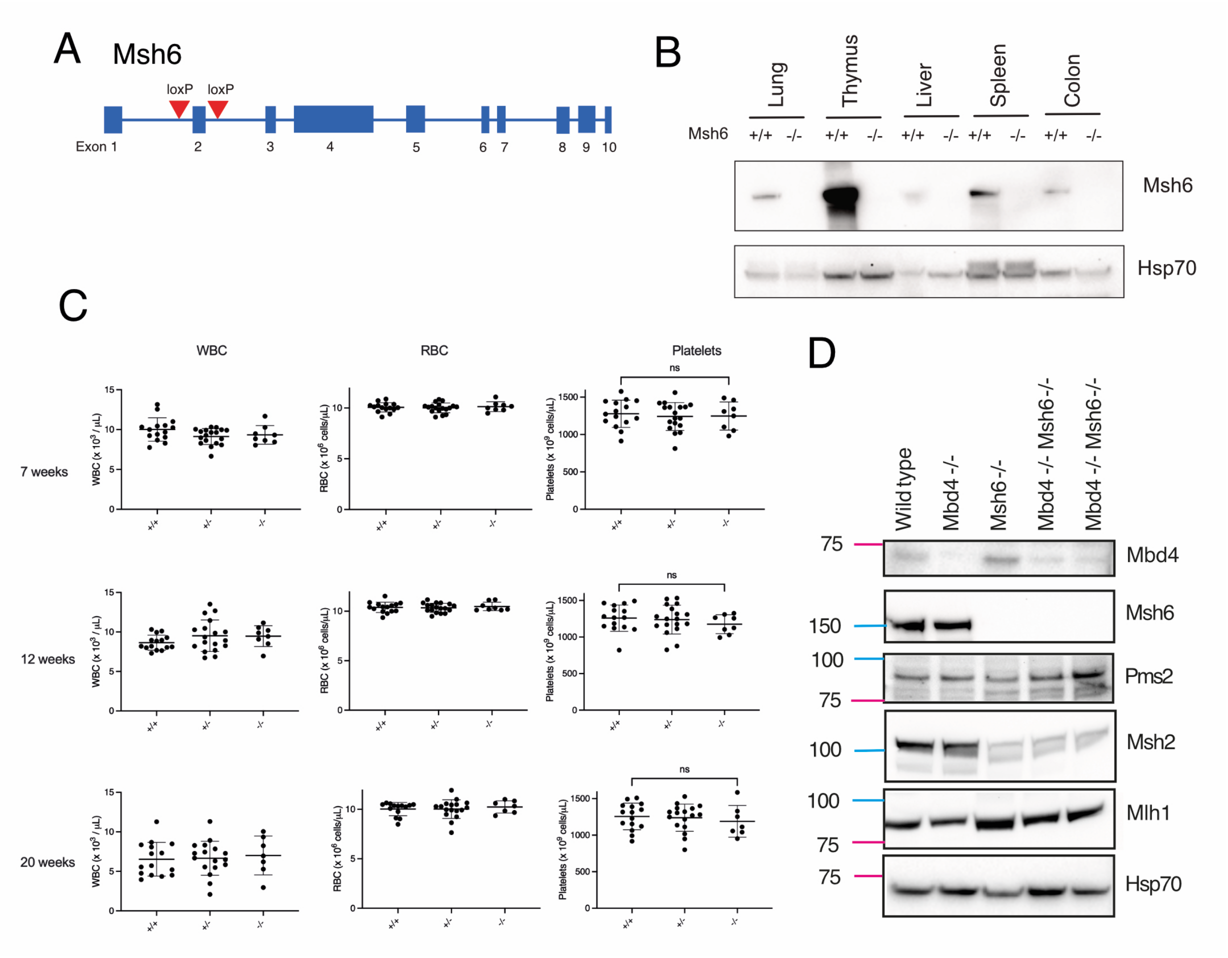


**Supplementary Figure 3: Characterisation of the *Msh6* knockout mouse model.** (A) Schematic of the floxed allele of *Msh6* that was used to generate the knockout model. Recombination at the flox sites surrounding exon 2 results in a premature stop codon. (B) Western blotting performed with lysates from mouse tissues demonstrating loss of the Msh6 protein in the knockout animals. Hsp70 was included as a loading control. (C) ADVIA analysis of peripheral blood in adult mice at different ages (7, 12 or 20 weeks). Counts are shown for WBC, RBC and platelets. Column labels reflect the *Msh6* genotype (+/+, +/- or -/-). No significant differences were observed (significance was tested by t-test with Welches correction). (D) Western blotting was performed with lysates from MEFs of the indicated genotype, with antibodies against the listed DNA repair proteins, anti-Mbd4 from R+D biosystems. Size markers are indicated at left (kDa).

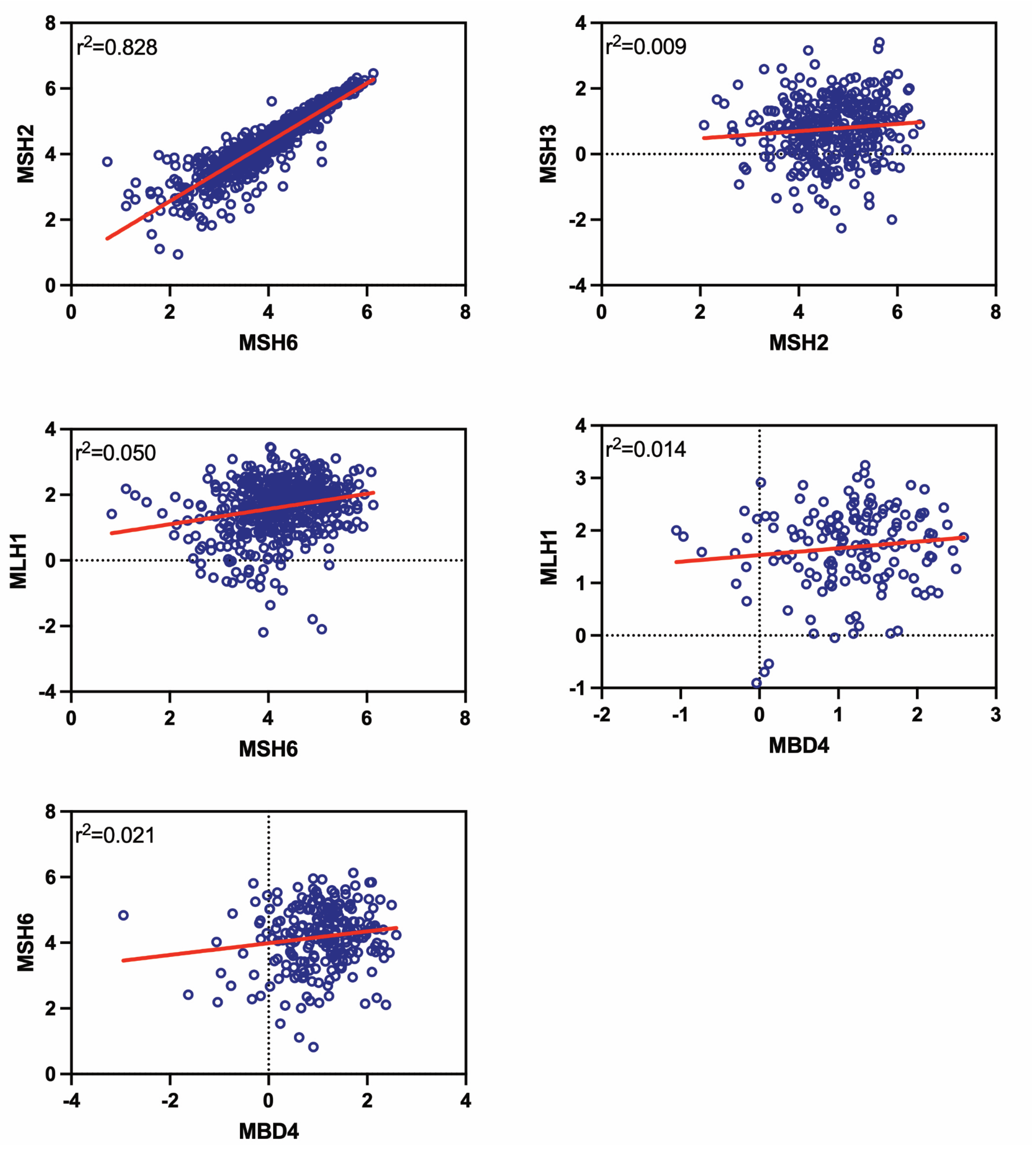


**Supplementary Figure 4: Correlation in DNA repair protein levels across human cancer cell lines.** Protein abundance estimates were taken from mass-spectrometry based profiling of human cancer cell lines (43). We assessed how protein expression was correlated between different MMR components and with MBD4. A linear regression line was fit and correlation scores ($r^2$) are shown.

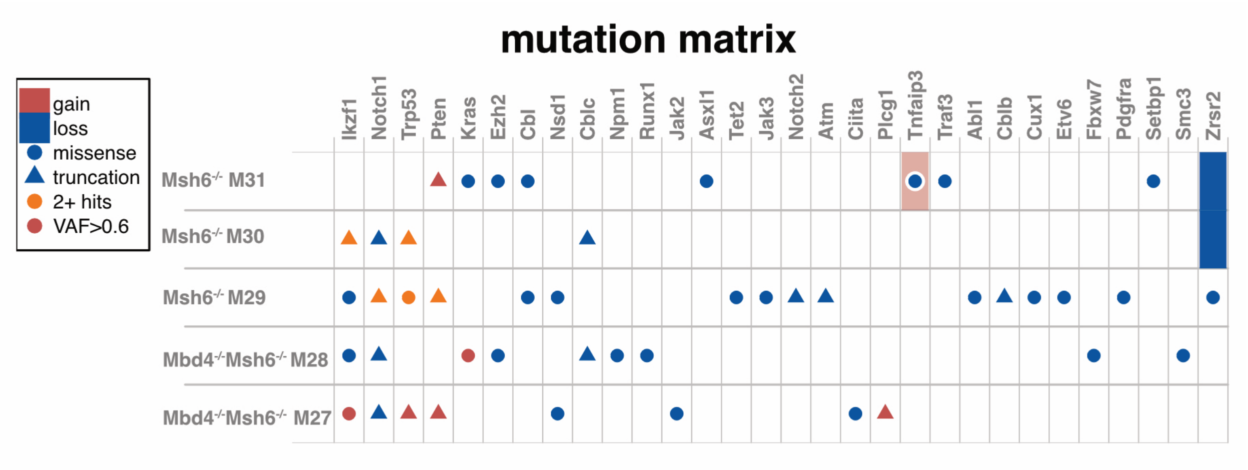


**Supplementary Figure 5: Driver mutation analysis of mouse thymomas.** A mutation matrix details candidate driver mutations in *Msh6*$^{-/-}$ and *Mbd4*$^{-/-}$*Msh6*$^{-/-}$ thymomas. Mutations were surveyed in genes with a high incidence of mutations in either myeloid or lymphoid cancers. Copy number estimates and point mutation status was derived with superFreq. Variant calls are provided as a supplementary dataset through Figshare.

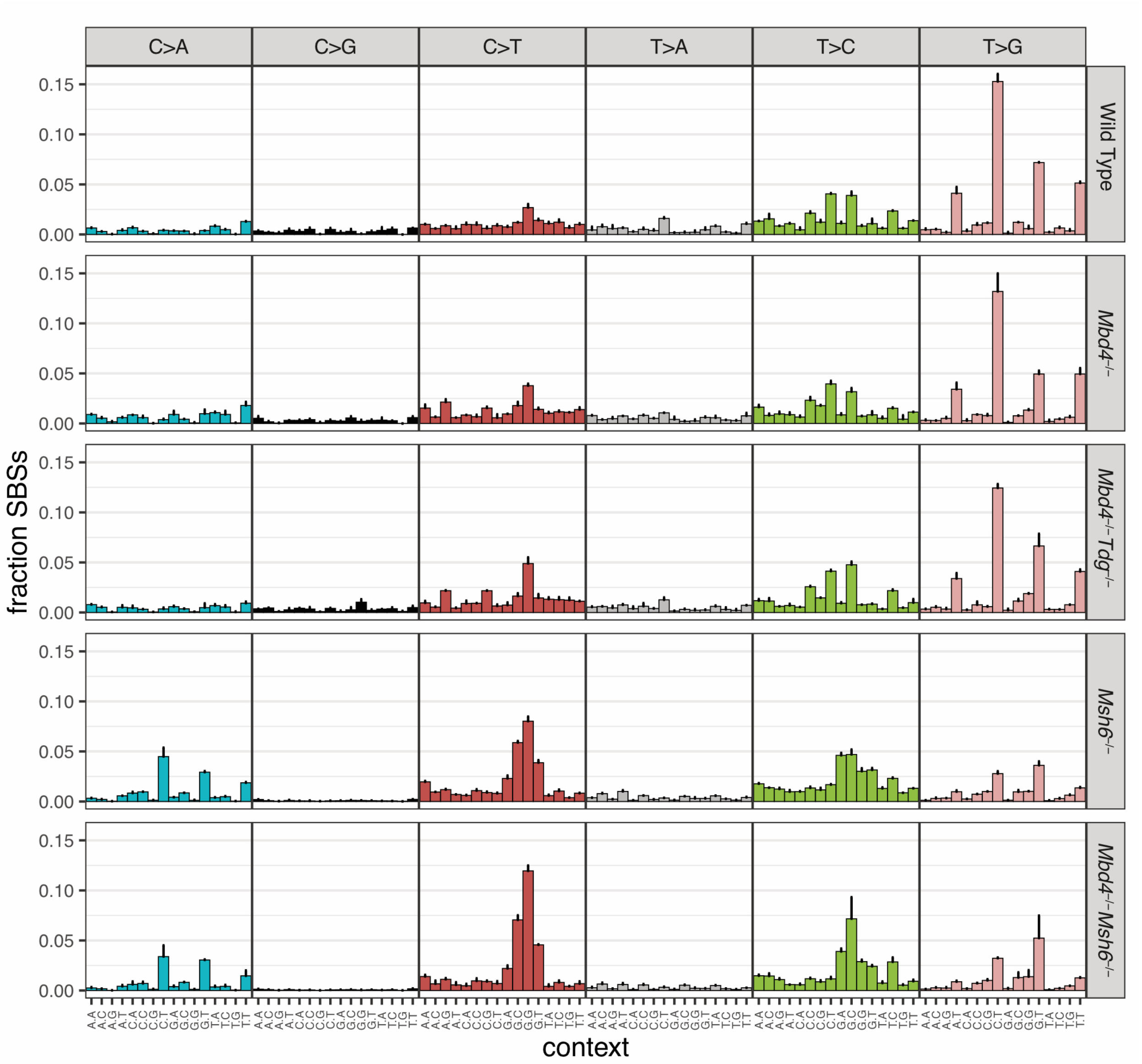


**Supplementary Figure 6: Mutational profiling in HoxA9-Meis1 myeloid cell lines.** Mutational signatures are shown as proportions for all long-term cultured myeloid cell lines. Values shown are mean ± SD. For each genotype we characterised 2 independent clones. Mutation rates (SBS/Mbp) for these cell lines are presented in Figure 1 and 3.

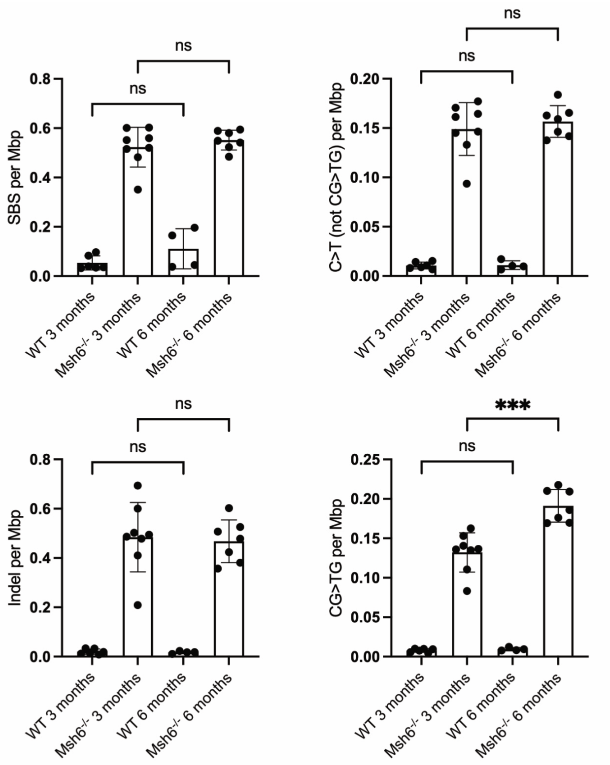


**Supplementary Figure 7: Assessment of mutation rate in aged bone marrow progenitors in *Msh6*$^{-/-}$ mice.** Somatic mutation rates were estimated for different classes of variants, either total SBS (top left), total indels (bottom left), C>T changes outside of CG sites (top right) or CG>TG mutations (bottom right). Each point represents an individual colony. Genotypes are shown below and data is separated based on the age of the mouse (either 3 months or 6 months). Error bars represent standard deviation and significance is based on unpaired t-test with Welch's correction.

**Supplementary Table Descriptions:**

**Supplementary Table 1:** Genotyping strategy and primers for mouse strains used in this study.

**Supplementary Table 2:** HoxA9-Meis1 cell line generation. CRISPR amplicon primer sequences and cell line genotypes for edits in *Mbd4* and *Msh6*.

**Supplementary Table 3:** Genotype outcomes for offspring from $Tdg^{+/-}$ intercrosses. Includes embryo genotyping at e10.5 and e12.5 and genotypes at weaning.

**Supplementary Table 4:** Summary of mutation rates across all genotypes and tissues for four classes of mutations, presented as mutations/Mb, or mutations/genome/day.

**Supplementary Table 5:** Coverage and descriptive statistics for all individual colonies, cell lines and tumour sequencing. Includes simulated and reference datasets. Includes sample labels, genotypes, age, tissue, coverage statistics, mutation counts, mutation rates, MMR signature decomposition, fraction methylated CG>TG, replication timing analysis and replication strand analysis.
